# Silaffins-Driven Genetic Engineering of Diatom Cell Walls: Insight into Biosilica Morphology and Nanomaterial Design

**DOI:** 10.1101/2024.12.20.629074

**Authors:** Tengsheng Qiao, Lulu Wang, Yan Zhao, Yun Li, Guanpin Yang, Baohua Zhu, Kehou Pan

## Abstract

Diatoms synthesize silica cell walls (frustules) with genetically encoded morphologies, ranging from nanopatterns to micropatterns, that far exceed current synthetic chemistry. Silaffins, a family of phosphoproteins undergoing complex post-translational modifications, have been isolated from frustules and shown to facilitate and regulate biosilica formation *in vitro* with long-chain polyamines. However, their particular role in frustule morphogenesis and functionality remains unclear. In this study, functions of two representative *silaffins*, *TpSil1* and *TpSil3*, were investigated in the model organism *Thalassiosira pseudonana* using gene overexpression and CRISPR/Cas9-mediated knockout approaches. Due to high sequence homology, *TpSil2* was concurrently disrupted in *TpSil1* knockout strains, while the homozygous knockout of *TpSil3* proved to be lethal. Quantitative morphological analysis revealed distinct yet complementary roles: TpSil3 governs both microscale overall size and mesoscale features, including macropore (fultoportula) density and mesopore (cribrum pore) pattern, whereas TpSil1/2 exclusively contribute to macropore morphogenesis and mesopore density. Overexpression of *silaffins* increased silica deposition, while knockouts exhibited reduced silicification but enhanced cell growth and photosynthetic efficiency. Furthermore, these genetic modifications significantly influenced the physicochemical and optical properties of bulk frustules, potentially enhancing the hemostatic, catalytic and photonic performances, thereby positioning them as versatile candidates for a wide range of biotechnological and industrial applications. Collectively, our findings elucidate the distinct roles of *TpSil1/2* and *TpSil3* in diatom physiology and frustule morphology, highlighting a promising pathway for engineering nanostructured silica materials with tailored properties through synthetic biology.

## 1. Introduction

Diatoms, with approximately 30,000 extant species (Mann and Vanormelingen, 2013), contribute about 20% to the global net biological primary production (Field et al., 1998). Among their most intriguing features is the biomineralized silica-based cell wall, known as the frustule. This structure is characterized by a highly ordered, three-dimensional porous architecture that spans from the nanoscale to the microscale (Kröger and Poulsen, 2008).

Frustules serve crucial function in diatom survival by providing protection against UV radiation (Aguirre et al., 2018), deterring grazers and parasites (Hamm et al., 2003), and facilitating essential physiological processes, including carbon acquisition (Milligan and Morel, 2002), nutrient uptake (Mitchell et al., 2013), and light scattering (Goessling et al., 2018). After diatom death, frustules remain in the environment as ballast in marine snow and fecal pellets, aiding in the vertical transport of carbon and silicon to the mesopelagic zone and deep ocean (Tréguer et al., 2018).

Beyond the ecological and physiological functions, frustules exhibit unique characteristics as innovative nanostructured biomaterials, including lightweight, high surface area, robust mechanical strength, thermal stability, biocompatibility, and the ability to modulate light. These attributes make frustules highly promising for various applications, including hemostasis (Wang et al., 2019), drug delivery (Delalat et al., 2015), enzyme immobilization ( Poulsen et al., 2007; Sheppard et al., 2012; Kumari et al., 2020), photonics (Ragni et al., 2018; Goessling et al., 2020) and so on. Moreover, the biosynthesis of frustules by diatoms is faithful, renewable, carbon neutral and highly efficient, surpassing the chemical synthesis methods. Elucidating the molecular mechanisms of frustule morphogenesis not merely advances our understanding of diatom physiology but also enables the eco-friendly, technological production of nanostructured silica materials with customizable properties through synthetic biology.

Frustules are composed of two interlocking halves, each consisting of a disk-shaped valve and several ring-like girdle bands (Mann, 1993). The morphogenesis of frustules is a complex, highly regulated process linked to the cell cycle, beginning with the uptake of Si(OH)4 by silicon transporter proteins in the plasma membrane (Shrestha and Hildebrand, 2015). Once internalized, Si(OH)4 is stored as soluble precursors and transported to the silica deposition vesicles (SDVs), where valves and girdle bands are synthesized. After silica precipitation, the structures are exocytosed from the SDVs and deposited onto the cell surface (Kröger, 2007; Hildebrand et al., 2018).

Diatoms consistently reproduce their species-specific frustules with remarkable precision across generations, underscoring the presence of a genetically encoded machinery that orchestrates the morphogenesis of their biosilica structures. The precise components of this machinery and their roles in driving biosilica morphogenesis remain key questions. Biochemical and proteomic analyses of isolated frustules and valve SDVs have identified over 100 proteins implicated in the morphogenetic process (Heintze et al., 2022; Kröger, 2022; Skeffington et al., 2022).

Transcriptomic studies of synchronized *Thalassiosira pseudonana* cultures also reveal a wide array of genes associated with frustule silicification (Mock et al., 2008; Shrestha et al., 2012; Brembu et al., 2017).

Over the past decade, functional genetic studies have identified several key proteins that regulate frustule morphology, including those involved in controlling overall frustule scale (TpSilacidin (Kirkham et al., 2017; Belshaw et al., 2023), silicalemma associated protein 2 in *Cyclotella cryptica* (CcSAP2, Wang et al., 2023)), the valve patterning center (SAP1 in *T. pseudonana* (TpSAP1, Tesson et al., 2017), valve z-height (TpSAP3 (Tesson et al., 2017), Silicanin-1 (TpSin1, Görlich et al., 2019)), as well as the number of fultoportulae (TpSilacidin, SAP2), and the density and area of cribrum pores (TpSSP20/THAPSDRAFT _21180 (Trofimov et al., 2019), TpdAnk1–3 (Heintze et al., 2022), TpSin1, SAP2). Moreover, genetic engineering has demonstrated the potential to enhance frustule properties. For instance, knockout of *TpSin1* resulted in frustules with enhanced catalytic activity of the immobilized enzyme on valves (Kumari et al., 2020), while *CcSAP2* knockdown frustules exhibited improved hemostatic performance (Wang et al., 2023). However, the preservation of frustule architectures in these mutants suggests that additional components are involved in frustule morphogenesis, underscoring the need for further exploration.

Among the proteins potentially involved in frustule morphogenesis, silaffins—zwitterionic phosphoproteins—are particularly intriguing. Named for their affinity with silica, silaffins are embedded within the silica matrix of the frustule, and can only be isolated using NH4F or anhydrous HF (Kröger et al., 1999; Poulsen and Kröger, 2004). Silaffins have been shown to facilitate silica polymerization *in vitro* through interaction with long-chain polyamines (LCPAs) (Kröger et al., 1999; Poulsen et al., 2003; Poulsen and Kröger, 2004) and have also been applied in biomimetic material fabrication (Abdelhamid and Pack, 2021).

To date, five silaffin genes have been identified: one from *Cylindrotheca fusiformis* (*sil1*) (Kröger et al., 2001) and four from *T. pseudonana* (*TpSil1*, *TpSil2*, *TpSil3* and *TpSil4*) (Poulsen and Kröger, 2004; Sumper and Brunner, 2008). These genes give rise to nine silaffin proteins through the proteolytic processing of their respective precursor polypeptides: natSil-1A1, natSil-1A2, natSil-1B, TpSil1H, TpSil1L, TpSil2H, TpSil2L, TpSil3 and TpSil4. However, the gene encoding natSil-2 of *C. fusiformis* has yet to be cloned due to the extensive posttranslational modifications (Poulsen et al., 2003). Notably, except for TpSil1 and TpSil2, silaffin polypeptides exhibit no significant sequence homology and are predicted to be intrinsically disordered proteins (Poulsen and Kröger, 2004; Scheffel et al., 2011). Despite this, they share similar amino acid compositions, unconventional sequence motifs, and extensive, diatom-specific, post-translational modifications (Poulsen and Kröger, 2004; Skeffington et al., 2022). Two main functional classes of silaffins have been proposed: i) “catalytic” silaffins, such as natSil-1A (Kröger et al., 2002) and TpSil1/2L (Poulsen and Kröger, 2004), which actively drive silica formation *in vitro*, and ii) “regulatory” silaffins, including natSil-2 (Poulsen et al., 2003), TpSil1/2H, and TpSil3 (Poulsen and Kröger, 2004), which modulate silica precipitation in a concentration-dependent manner, promoting the formation of flat plates or porous silica that resemble the frustule. The regulatory silaffins likely fine-tune silica deposition *in vivo*, directing it to specific regions and facilitating the formation of intricate, species-specific morphologies. Despite significant advances in the structural and *in vitro* functional characterizations of silaffins, much remains to be uncovered regarding how their roles in frustule morphogenesis and functionality.

*T. pseudonana*, the first diatom species to have its genome sequenced (Armbrust et al., 2004), has become a model organism for studying frustule morphogenesis due to its global distribution, simple frustule architecture, and genetic modifiability (Poulsen et al., 2006; Hopes et al., 2016; Poulsen and Kröger, 2023). Four silaffin genes have been identified in *T. pseudonana*. *TpSil1* and *TpSil2* share a remarkably high sequence identity (93.56%, SI Appendix, Fig. S2), with only one corresponding gene THAPSDRAFT_11360 present in the genome. PCR amplification and DNA sequencing revealed identical promoter sequences for both genes (data not shown), suggesting they probably function as allelic variants. In terms of subcellular localization, the TpSil1-GFP fusion protein localizes specifically around the fultoportulae of the valve, whereas TpSil3-GFP is broadly distributed across the entire frustule. The localization of TpSil4-GFP resembles that of Sil3-GFP (Poulsen et al., 2013); however, the transcriptional abundance of *TpSil4* is approximately 50 times lower than *TpSil3* (data not shown). Furthermore, the expression patterns of *TpSil1* and *TpSil3* are tightly synchronized with specific stages of frustule synthesis (Frigeri et al., 2006; Hildebrand et al., 2007). Based on these findings, this study focuses on *TpSil1* and *TpSil3* as representative *silaffin* genes. Their roles in diatom physiological processes (cell growth, photosynthesis and silicification), individual frustule morphologies (valve size, fultoportula and cribrum pore patterns) and bulk frustule properties (BET surface area, particle size distribution, chemical structure, surface wettability and optical properties) were investigated through gene overexpression and CRISPR/Cas9-mediated knockout approaches.

## 2. Materials and methods

### 2.1 Strains and Growth Conditions

The wild-type (WT) and genetically transformed strains of *Thalassiosira pseudonana* CCMP 1335 (Laboratory of Applied Microalgae Biology, Ocean University of China) were grown in Aquil medium at 20℃ under continuous light (120-140 μmol photons m^-2^ s^-1^). The composition of Aquil medium is available on the Bigalow website. The nitrate concentration of 550 µM and phosphate concentration of 55 µM were used in this study.

Before biolistic transformation, *T. pseudonana* was pre-cultured in half-salinity Aquil medium to optimize nourseothricin selection (Hopes et al., 2017). Phenotypic analyses were conducted on WT and genetically transformed strains after culturing in Aquil medium for more than three months.

### 2.2 Plasmid Constructions

The pPha-T1 vector (Zaslavskaia et al., 2000) was used as the backbone. The coding sequences of silaffins or Cas9 were placed under the nitrate reductase (*NR*) cassette. Single guide RNAs (sgRNAs) targeting *TpSil1* and *TpSil3* were designed using CRISPOR (Concordet and Haeussler, 2018) and CRISPRdirect (Naito et al., 2015). DNA fragments of potential off-target sites for mismatched sgRNAs were not analyzed, as previous studies showed that sites with four or more mismatches were unaffected in *Phaeodactylum tricornutum* (Stukenberg et al., 2018) and *T. pseudonana* (Görlich et al., 2019). Two sgRNAs per target gene were each placed under the *T. pseudonana* U6 promoter (Hopes et al., 2016). Details on the constructions of the nourseothricin resistance, *silaffin* gene overexpression and knockout plasmids, as well as the sgRNA design, *in vitro* sgRNA efficacy validation and off-target prediction, are provided in SI Appendix (Supplementary Text S1).

### 2.3 Biolistic Transformation

The plasmids were introduced into *T. pseudonana* cells using the PDS-1000/He particle delivery system (Bio-Rad) as described previously ( Poulsen et al., 2006; Hopes et al., 2017). Briefly, *T. pseudonana* cells in the exponential growth phase were centrifuged at 3,000 g for 10 min, and approximately 1 × 10^8^ cells were plated onto the center of plates (1.5% agar, half-salinity). Plasmid DNA for transformation was extracted using the E.Z.N.A.^®^ Endo-free Plasmid Kit (OMEGA, D6950). Biolistic transformation was performed by bombarding tungsten particles (M-10, 0.7 µm diameter, Bio-Rad) coated with approximately 10 μg plasmid DNA into *T. pseudonana* cells under vacuum at 1350 psi from a distance of 6 cm. Following bombardment, cells were transferred into liquid medium at approximately 1 × 10^6^ cells mL^-1^ and incubated under constant light for 24 h. The transformed cells were then centrifuged at 3,000 g for 10 min and plated approximately 2 × 10^7^ cells per plate (0.8% agar, half-salinity) containing 100 µg mL^-1^ nourseothricin (Jena Bioscience). Resistant colonies appeared after 8–12 days and were transferred to liquid medium containing 100 µg mL^-1^ nourseothricin. For re-plating, approximately 1 × 10^3^ cells were plated onto plates (0.8% agar, half-salinity) containing 100 µg mL^-1^ nourseothricin.

### 2.4 Verification of Genetically Transformed Strains

For the genomic screening of the genetically transformed strains, details are provided in SI Appendix (Supplementary Text S2). For the genetic modified strains, overexpression of *TpSil1* strains are denoted as OE1-1 and OE1-2; *TpSil3* overexpression as OE3-1 and OE3-2; *TpSil1* and *TpSil2* double knockout strains are labeled as KO1/2-1 and KO1/2-2; and the incomplete knockout of *TpSil3* is represented as iKO3-1 and iKO3-2.

For the transcript verification, the WT and genetically transformed strains of *T. pseudonana* cultured to the logarithmic growth phase were collected and stored at -80℃. Total RNA was extracted from the cultures using the E.Z.N.A.^®^ Total RNA Kit II (OMEGA, R6934). Approximately 100 ng of RNA was reverse-transcribed into cDNA using the HiScript III RT SuperMix (Vazyme, R323). The relative gene expression levels were quantified using the Applied Biosystems QuantStudio5 Real-Time PCR System. The actin-like protein gene (protein ID 269504) (Durkin et al., 2009) was used as a reference gene. The 2^-ΔΔCT^ method (Livak and Schmittgen, 2001) was applied to calculate the relative expression levels of the target genes. The qRT-PCR primer sequences are listed in SI Appendix (Table S2).

### 2.5 Diatom Growth and Photosynthetic Efficiency Measurement

The growth performance of diatom was evaluated by cell counting using hemocytometers. Growth rate was determined using the formula: (N1 − N0) ⁄ (t1 − t0), where N1 and N0 represent the cell densities at times t1 and t0, respectively.

To evaluate the photosynthetic performance, chlorophyll fluorescence parameters of photosystem II (PSII), including the maximum quantum yield (*Fv*/*Fm*), effective quantum yield (YII) and electron transport rate (ETR), were monitored using a PAM fluorometer (Water-PAM, Walz). Prior to measurement, cells were dark-acclimated for 15 min.

### 2.6 Cellular Silica Content Determination

Silica content was quantified using the molybdate blue assay (Ramachandran and Gupta, 1985), following protocols reported by Wang et al., (2023). Briefly, 1 × 10^7^ cells (exact counts recorded) were centrifuged at 3,000 g for 10 min, washed once with 2 mL of methanol, followed by two washes with 5 mL of distilled water, and resuspended in 1 mL of water. Next, 500 μL of 6 M NaOH was added, and the mixture was incubated at 95℃ for 1 h. To measure silicic acid content, 50 μL of the resulting supernatant was mixed with 100 μL of distilled water and 50 μL of molybdate solution. After incubation at 25℃ for 10 min, 75 μL of reducing agent was added, and the mixture was further incubated at 25℃ for 3 h. Absorbance at 810 nm was measured, and the silica content was determined using a sodium hexafluorosilicate standard curve. Silica content per cell was calculated by normalizing the concentrations to the recorded cell counts.

### 2.7 Frustule Extraction

The preparation of diatom frustules for scanning electron microscopy (SEM) followed established protocols (Görlich et al., 2019). Briefly, approximately 1 × 10^8^ cells were collected by centrifugation at 3,000 g for 10 min. The cell pellets were washed three times with distilled water, and resuspended in 15 mL of extraction buffer (2% SDS, 100 mM EDTA, pH 8, 1 mM PMSF). The suspension was incubated at 60℃ with continuous shaking for 1 h, then pelleted by centrifugation (3,000 g, 2 min) and washed three times with 15 mL of washing buffer (10 mM EDTA, pH 8, 1 mM PMSF). This extraction and washing process was repeated until the frustules became colorless. Subsequently, the frustules were washed with 2 mL of acetone, followed by four washes with 15 mL of distilled water, and finally resuspended in 200 µL water.

For physicochemical and optical property analyses, bulk frustules were prepared as described in by Wang et al., (2019). Approximately 1 × 10^11^ cells were harvested by centrifugation at 3,000 g for 10 min. The cell pellets were washed three times with distilled water and extracted with 50 mL of sulfuric acid (98%) at 60℃ for 1 h, followed by incubation at room temperature for 48 h. Next, 50 mL of nitric acid (65%) was added for further extraction at 60℃ for 4 h. After cooling to room temperature, the frustules were collected by centrifugation (3,000 g, 5 min) and subsequently washed with 5 mL of acetone, four times with distilled water, and resuspended in 2 mL of water. The extracted frustules were frozen at -80℃ and freeze-dried using a vacuum freeze dryer. Genetically transformed strains OE1-1, OE3-1, KO1/2-1, iKO3-1 were selected for bulk frustule property analysis.

### 2.8 Imaging and Morphological Characteristics Quantitative Determination

For light microscopy analysis, cells were observed using a Nikon Eclipse 80i optical microscope. Given the cylindrical shape of diatom cells, which appear as either circular or cylindrical profiles under the microscope, statistical analyses were performed on 30 randomly selected cells per strain for each profile type. Cell dimensions were determined by calculating cell areas in the images using ImageJ (SI Appendix, Fig. S13A). Specifically, images were converted to grayscale and thresholded to isolate the cell area, followed by manual selection and measurement using the polygon or oval tool. The scale was calibrated based on the scale bar provided in the images.

Frustules were gold-coated using a sputter coater and imaged with a Zeiss GeminiSEM 360 scanning electron microscope. To capture detailed morphological characteristics, 12 frustules per strain were analyzed using ImageJ and Python 3.9 libraries. Valve diameter and the number of fultoportulae were measured in ImageJ, with calibration performed using the scale bar in the SEM images (SI Appendix, Fig. S13B). SEM images were segmented into multiple 60 × 60-pixel sub-images (333.33 × 333.33 nm per sub-image). For fultoportulae mean area measurements, sub-images specifically focusing on fultoportulae regions were selected, and areas were calculated from two front-facing fultoportulae per frustule (SI Appendix, Fig. S14A). For cribrum pore pattern analysis, sub-images were exclusively focused on pore regions, with six sub-images analyzed per frustule. To enhance pore clarity, GaussianBlur from OpenCV, with a 5×5 kernel, was applied to reduce noise, followed by histogram equalization (cv2.equalizeHist) to improve contrast and sharpen edges, facilitating precise annotation of pore boundaries. The areas of fultoportulae and cribrum pores were manually annotated using LabelMe. The resulting data, stored in JSON format, were processed to generate binary masks using ImageDraw.Draw, where pore regions were represented in white (255) and the background in black (0). Subsequently, the skimage.measure.label function and regionprops were employed to extract geometric features. Finally, pixel-based measurements were converted to physical units according to the image scale bar (SI Appendix, Fig. S14B). The pore-to-total-area ratio was calculated by normalizing the total pore area to the corresponding sub-image area.

### 2.9 Frustule Structural and Optical Properties Analysis

To evaluate the total surface area, pore volume, and pore diameter of bulk frustules, Brunauer-Emmett-Teller (BET) analysis was conducted with a Micromeritics 3Flex Surface Characterization Analyzer. Approximately 100 mg of frustules were degassed at room temperature for 12 h prior to nitrogen adsorption measurements. Contact angle (CA) measurements were performed using distilled water with a Dataphysics OCA20 goniometer to reveal surface wettability. The particle size distribution of the frustules was analyzed using a Malvern Mastersizer 3000 laser diffraction analyzer with ethanol as the dispersing agent. The resulting data were analyzed using Mastersizer 3000 software (version 3.88). Fourier transform infrared spectroscopy (FTIR) was performed using a Thermo Scientific NICOLET 6700 FTIR spectrometer, in which the frustule samples were prepared on potassium bromide (KBr) disks and measured in ambient air. UV-visible diffuse reflectance spectroscopy (UV-Vis DRS) was conducted with a Shimadzu UV-3600i Plus spectrophotometer to analyze the optical absorbance and reflectance properties of the frustules. Photoluminescence (PL) spectra were recorded with an Edinburgh Instruments FLS980 spectrometer to evaluate the luminescent properties under excitation wavelengths of 330 and 380 nm.

### 2.10 Statistical Analysis

The number of replicates for each strain is provided in the respective figure and table legend. Statistical comparisons between two groups were performed using the Python SciPy library with an independent two-sample t-test, assuming equal variance. *P*-values were calculated to evaluate statistical significance between different strains. Asterisks indicate significance levels (\**P* < 0.05, \*\**P* < 0.01, \*\*\**P* < 0.001), with "ns" denoting non-significant differences. Black lines are drawn to connect the compared groups where applicable.

## 3. Results

### 3.1 Silaffins Gene Overexpression and Knockout Strains

Genomic characterization of *TpSil1* and *TpSil3* overexpression strains of *T. pseudonana* is detailed in SI Appendix (Supplementary Text S2.1). Two strains for each gene (OE1-1 and OE1-2 for *TpSil1*, OE3-1 and OE3-2 for *TpSil3*) were selected for transcriptional validation. Due to the high sequence homology between *TpSil1* and *TpSil2*, qRT-PCR primers designed for *TpSil1* also targeted transcripts from *TpSil2*. qRT-PCR analysis revealed a significant upregulation of *TpSil1/2* expression in both OE1-1 and OE1-2 strains, with expression levels 3.29- and 2.73-fold higher than the WT, confirming successful *TpSil1* overexpression at the transcriptional level (Fig. 1B). *TpSil3* expression levels were markedly upregulated in OE3-1 and OE3-2 strains, showing 16.47- and 21.52-fold higher than WT, indicating robust *TpSil3* transcriptional overexpression (Fig. 1C).

**Fig. 1.**
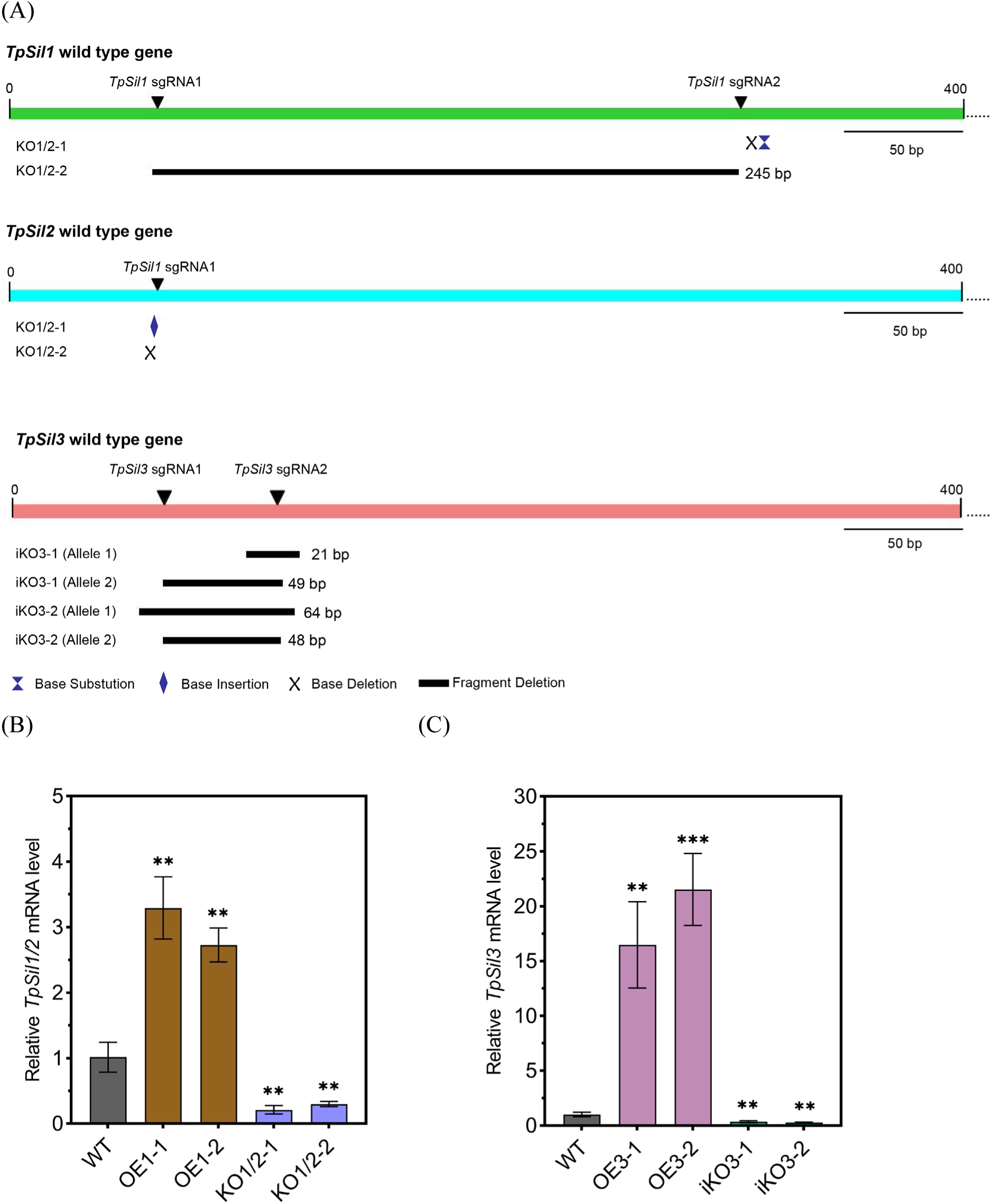
Characterization of the *silaffin* overexpression and knockout strains of *T. pseudonana*. (A) Bar graphs depicting segments of each *silaffin* gene (first 400 base pairs from the 5’ end), with vertical arrows indicating sgRNA binding sites and highlighted edited regions for each knockout strain. (B) Relative mRNA expression levels of *TpSil1/2*, and (C) Relative mRNA expression levels of *TpSil3*, normalized to the actin-like protein reference gene (ID 269504). Strains: WT: wild-type strain; OE1-1 and OE1-2: *TpSil1* overexpression strains; KO1/2-1 and KO1/2-2: *TpSil1* and *TpSil2* double-knockout strains; OE3-1 and OE3-2: *TpSil3* overexpression strains; iKO3-1 and iKO3-2: incomplete *TpSil3* knockout strains. Data are presented as mean ± standard deviation (SD), with *n* = 3 biologically independent samples. Statistical comparisons were performed to the wild-type strain using an independent two-sample t-test with equal variance assumptions (\**P* < 0.05, \*\**P* < 0.01, \*\*\**P* < 0.001).

Screening of *TpSil1* knockout strains is described in SI Appendix (Supplementary Text S2.2.1). Due to the high sequence similarity between *TpSil1* and *TpSil2*, the *TpSil1_*sgRNA1, designed to specifically target *TpSil1*, also targeted *TpSil2* (SI Appendix, Fig. S4 and Table S3). DNA sequencing confirmed the knockout of both *TpSil1* and *TpSil2* sequences, resulting in the disruption of both genes. The edited regions in these strains are indicated in Fig. 1A. Sequencing results for KO1/2-1 and KO1/2-2 (SI Appendix, Fig. S11A) confirmed that the disrupted *TpSil1* and *TpSil2* produce significantly truncated polypeptides (SI Appendix, Fig. S12). Relative expression levels of *TpSil1/2* were significantly downregulated in KO1/2-1 (by 79.20%) and KO1/2-2 (by 70.52%) (Fig. 1B).

Screening of *TpSil3* knockout strains was challenging (SI Appendix, Supplementary Text S2.2.2). Among the 168 identified strains, each retained at least one *TpSil3* allele with either its full sequence intact or a base deletion in multiples of three (SI Appendix, Fig. S11B), resulting in no deletion or only a small number of deleted amino acids (SI Appendix, Fig. S12), indicating that homozygous knockout of *TpSil3* is probably lethal. Consequently, two incomplete *TpSil3* knockout strains, iKO3-1 and iKO3-2, were identified and chosen for further investigation. The edited regions in these strains are shown in Fig. 1A. Relative *TpSil3* expression levels were significantly downregulated in iKO3-1 (by 64.15%) and iKO3-2 (by 71.43%) (Fig. 1C).

Protein-level validation via western blot (WB) analysis was not feasible for TpSil1 and TpSil3. Both TpSil1-GFP and TpSil3-GFP fusion proteins were tightly bound to the frustules, and treatment with 1% SDS, a standard protein extraction detergent, failed to release the GFP-tagged silaffins from frustules (Poulsen et al., 2013), suggesting their entrapment within the frustule’s silica matrix. Subsequent phenotype analysis was performed on WT and genetically transformed strains of *T. pseudonana* that had been autotrophically cultured for over three months to eliminate the effects of biolistic bombardment.

### 3.2 Silaffins on Physiological Characteristic of Diatom

Growth rate, photosynthetic efficiency, and silica content were assessed to evaluate the role of silaffins in diatom physiology. Growth rate analysis (Fig. 2A) showed that the WT strain of *T. pseudonana* grew at a rate of (5.75 ± 0.22) × 10^5^ cells mL^-1^ d^-1^ during 9 days of cultivation. The OE1, OE3, and KO1/2 strains exhibited growth rates comparable to WT. Notably, iKO3 demonstrated a significantly elevated growth rate (*P* < 0.01) of (6.37 ± 0.18) × 10^5^ cells mL^-1^ d^-1^, indicating that *TpSil3* disruption may enhance protoplast proliferation.

**Fig. 2.**
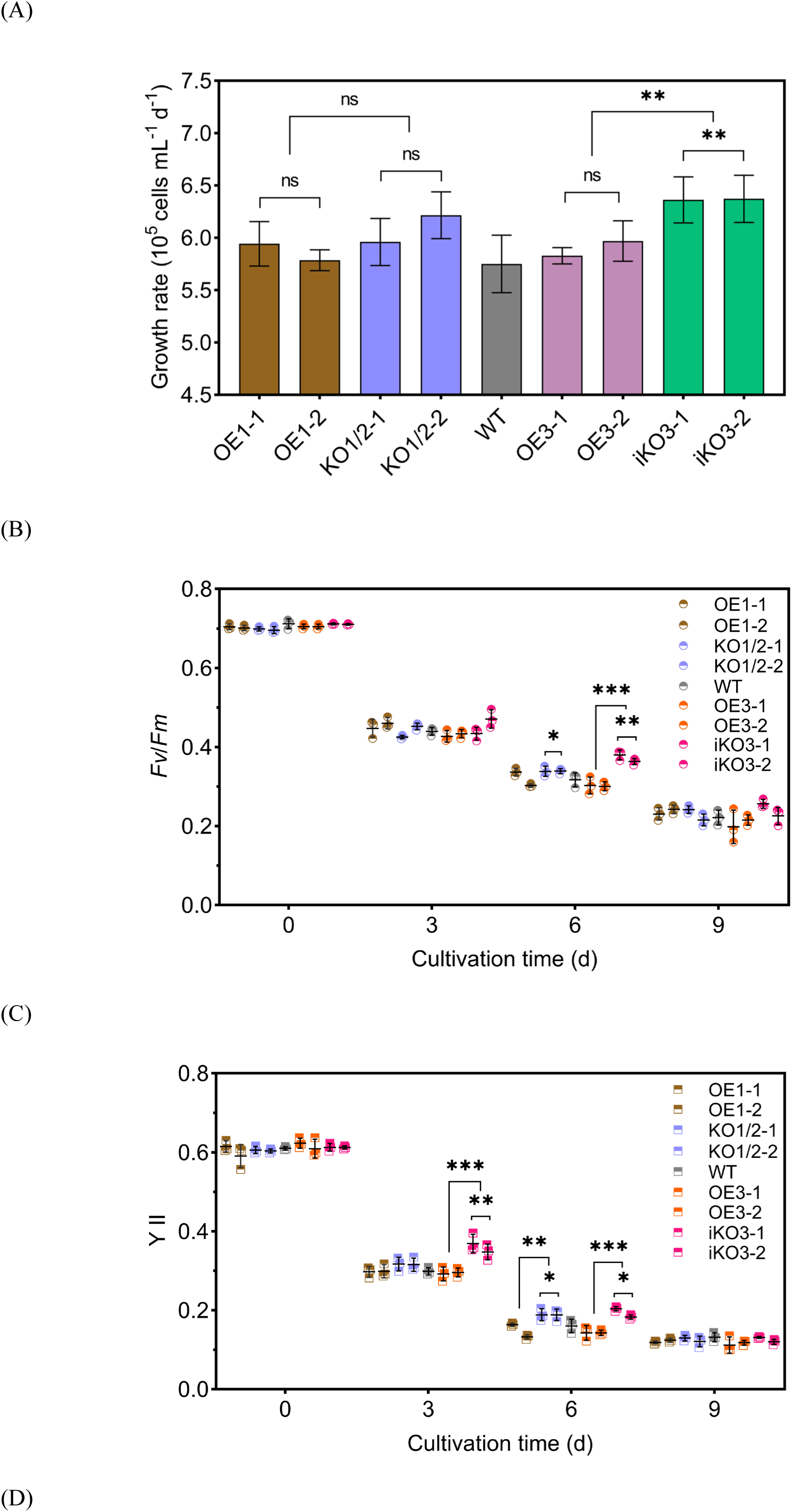
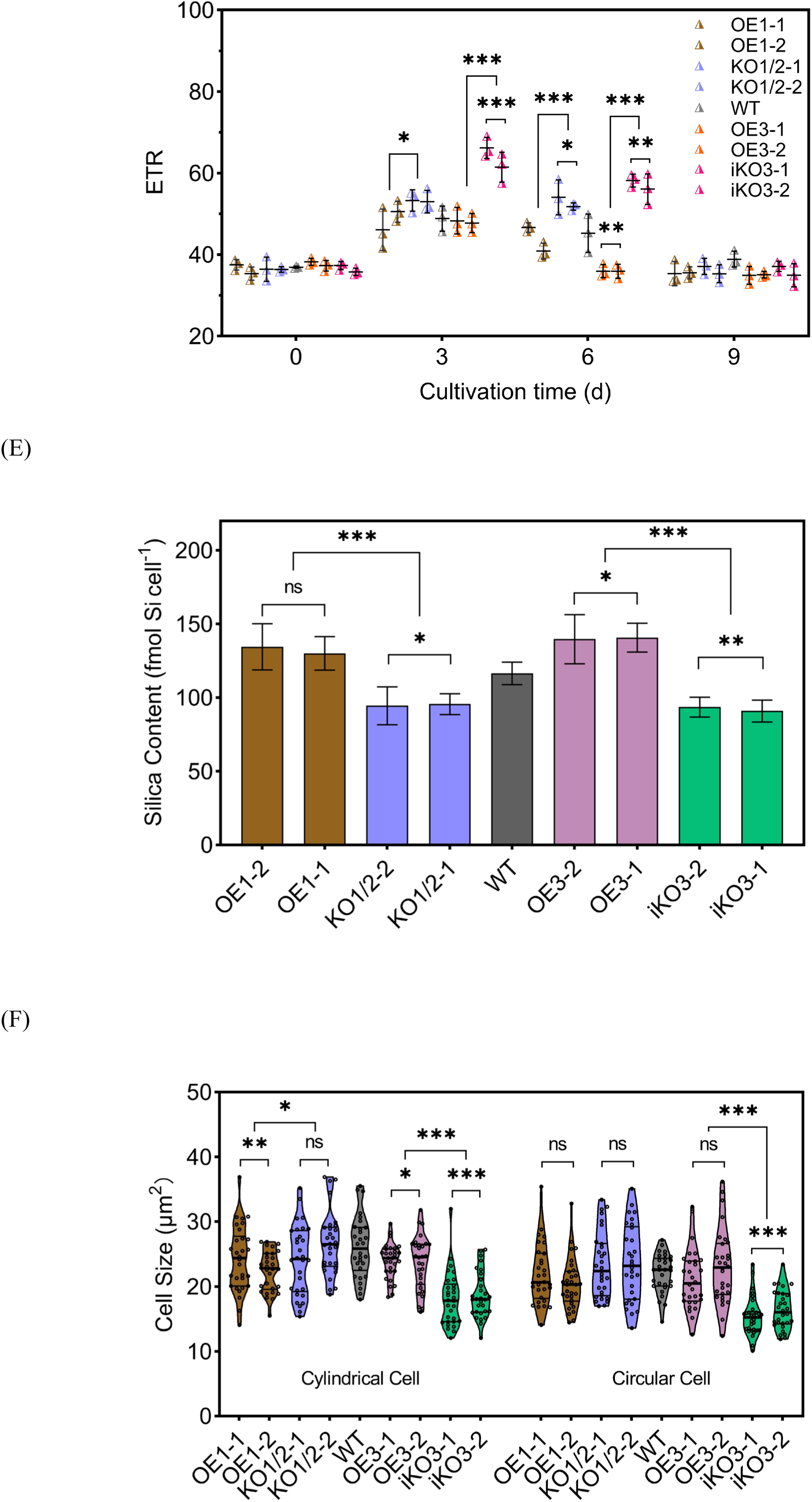
Growth rates, photosynthetic performance, cellular silica content, and cell dimensions of wild-type and genetically transformed *T. pseudonana* strains. (A) Growth rate measured after 9 days of cultivation. (B) Maximum quantum yield of PSII (*Fv/Fm*). (C) Effective quantum yield of PSII (YII). (D) Electron transport rate (ETR). (E) Cellular silica content. Data are presented as mean ± standard deviation (SD), with *n* = 3 biologically independent samples. (F) Cell surface area observed by light microscopy. 30 cells per strain were measured for each profile type (*n* = 30). Thin lines represent the quartiles, the thick line marks the median, and individual data points are displayed as dots within the violin plots. Strains: WT: wild-type strain; OE1-1 and OE1-2: *TpSil1* overexpression strains; KO1/2-1 and KO1/2-2: *TpSil1* and *TpSil2* double-knockout strains; OE3-1 and OE3-2: *TpSil3* overexpression strains. iKO3-1 and iKO3-2: incomplete *TpSil3* knockout strains. Statistical comparisons between two groups were performed using an independent two-sample t-test with equal variance assumptions (\**P* < 0.05, \*\**P* < 0.01, \*\*\**P* < 0.001).

Across the 9-day cultivation, the maximum quantum yield of photosystem II (*Fv*/*Fm*) remained consistent in WT, OE1, and OE3 strains (Fig. 2B). However, KO1/2 strains exhibited significantly higher *Fv*/*Fm* on day 6 (*P* < 0.05), with iKO3 strains showing an even greater increase (*P* < 0.01). The effective quantum yield (YII, Fig. 2C) followed a similar pattern, with iKO3 strains showing a significant increase by day 3 (*P* < 0.01). For the electron transport rate (ETR, Fig. 2D), OE3 strains displayed lower ETR on day 6 (*P* < 0.01), while KO1 and iKO3 strains exhibited higher values than WT on day 6 (*P* < 0.05 and *P* < 0.001, respectively). These results suggest enhanced photosynthetic efficiency in *silaffin* knockout strains, particularly in iKO3.

In terms of cellular silica content, (Fig. 2E), which reflects both intracellular and frustule silica levels, OE1 strains exhibited slight, statistically insignificant increases compared to WT. KO1/2 strains had significantly lower silica content (*P* < 0.001 compared to OE1, *P* < 0.05 compared to WT). Similarly, OE3 strains displayed marked increases in silica content (*P* < 0.05), while *TpSil3* knockout strains demonstrated significantly lower levels than WT (*P* < 0.01). These findings imply that TpSil1/2 and TpSil3 play pivotal roles in diatom silicification, potentially through direct interaction with silica precursors or interplay with other regulatory proteins.

To investigate the role of *silaffins* in diatom morphology, cell size was measured using light microscopy (Fig. 2F, SI Appendix, Fig. S13A). Due to the cylindrical shape of diatom cells, both circular and cylindrical profiles were observed. But measurements based on the circular cell profile were more accurate as the cylindrical profile can be influenced by the cell division state. The WT strain had an average cylindrical profile area of 26.09 ± 4.59 µm^2^ and a circular profile area of 22.10 ± 2.89 µm^2^. No significant differences were observed in circular profile dimensions between WT and OE1, OE3, or KO1/2 strains. However, iKO3 strains exhibited a significant reduction for both cell profiles (*P* < 0.001), indicating that *TpSil3* is essential for maintaining cell size. Further investigation into frustule morphology in *silaffin* mutants will offer deeper insights into the structural and functional roles of silaffins in diatom silicification.

### 3.3 Silaffins on Morphological Characteristic of Frustule Valve

The frustule valve of *T. pseudonana* features a circular arrangement of radially branching silica ribs, termed costae (green lines in Fig. 3E), interconnected by cross-connections (orange lines in Fig. 3E) that form trapezoidal areola pores (red rectangles in Fig. 3E). Each areola pore contains small cribrum pores (blue dots in Fig. 3E). Fultoportulae (yellow circles in Fig. 3E), symmetrically arranged near the valve margin with one or two centrally positioned, consist of a central tube surrounded by 2–4 satellite pores (Babenko et al., 2022). The girdle bands have smooth surfaces with regular pores. The intricate valve patterns are the primary focus in studies of frustule morphology. SEM analysis revealed consistent circular valve shapes and ridge networks across all valves (Fig. 3). The valve characteristics of OE1 and OE3 were found closely resemble those of the WT. However, KO1/2 frustules showed reduced cross-connections, thickened costae, and exposed satellite pores of fultoportulae (Fig. 3C, D). iKO3 frustules exhibited significant morphological changes, including smaller valve size, fewer fultoportulae, and reduced costae and cross-connections (Fig. 3I, J).

**Fig. 3.**
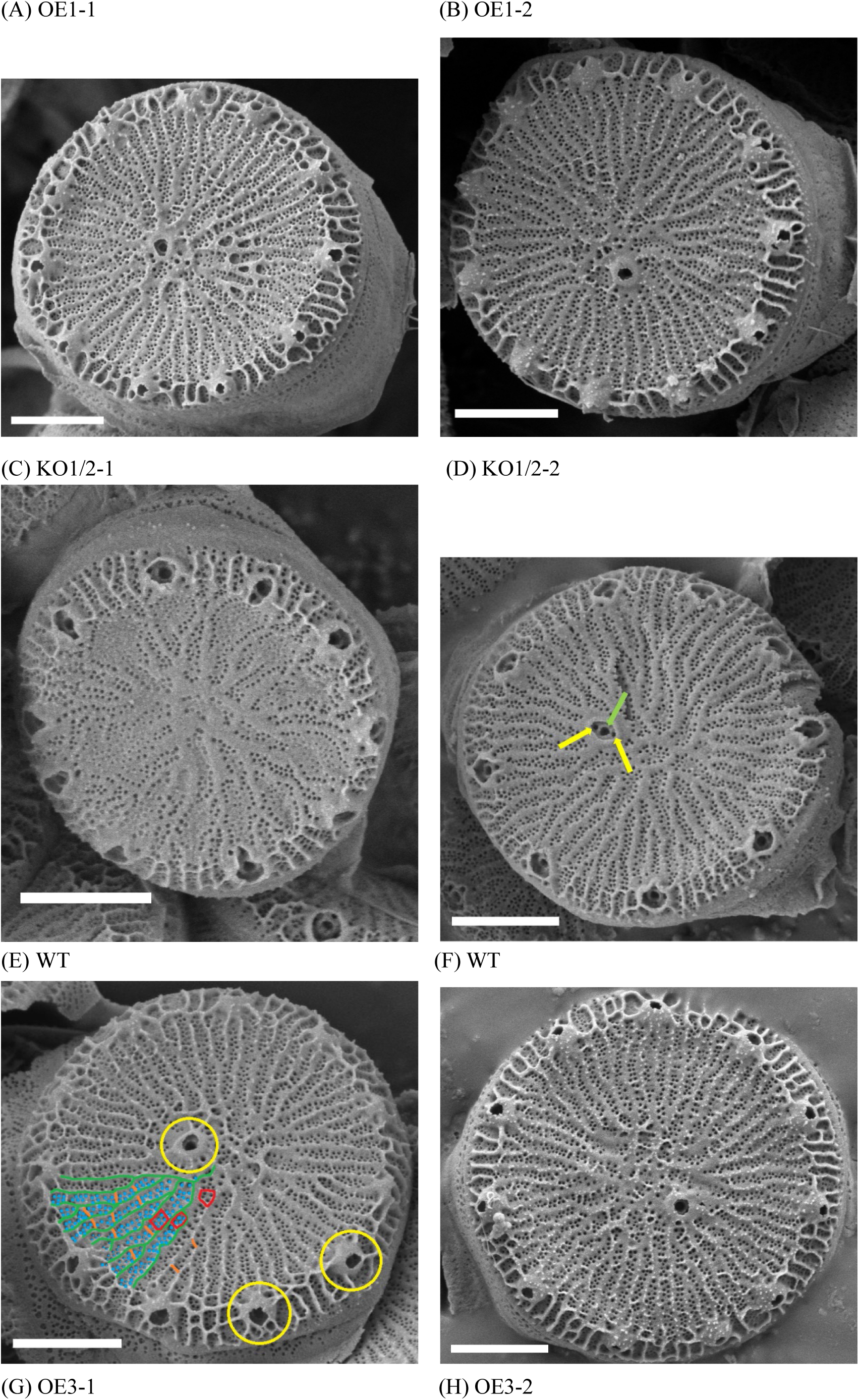
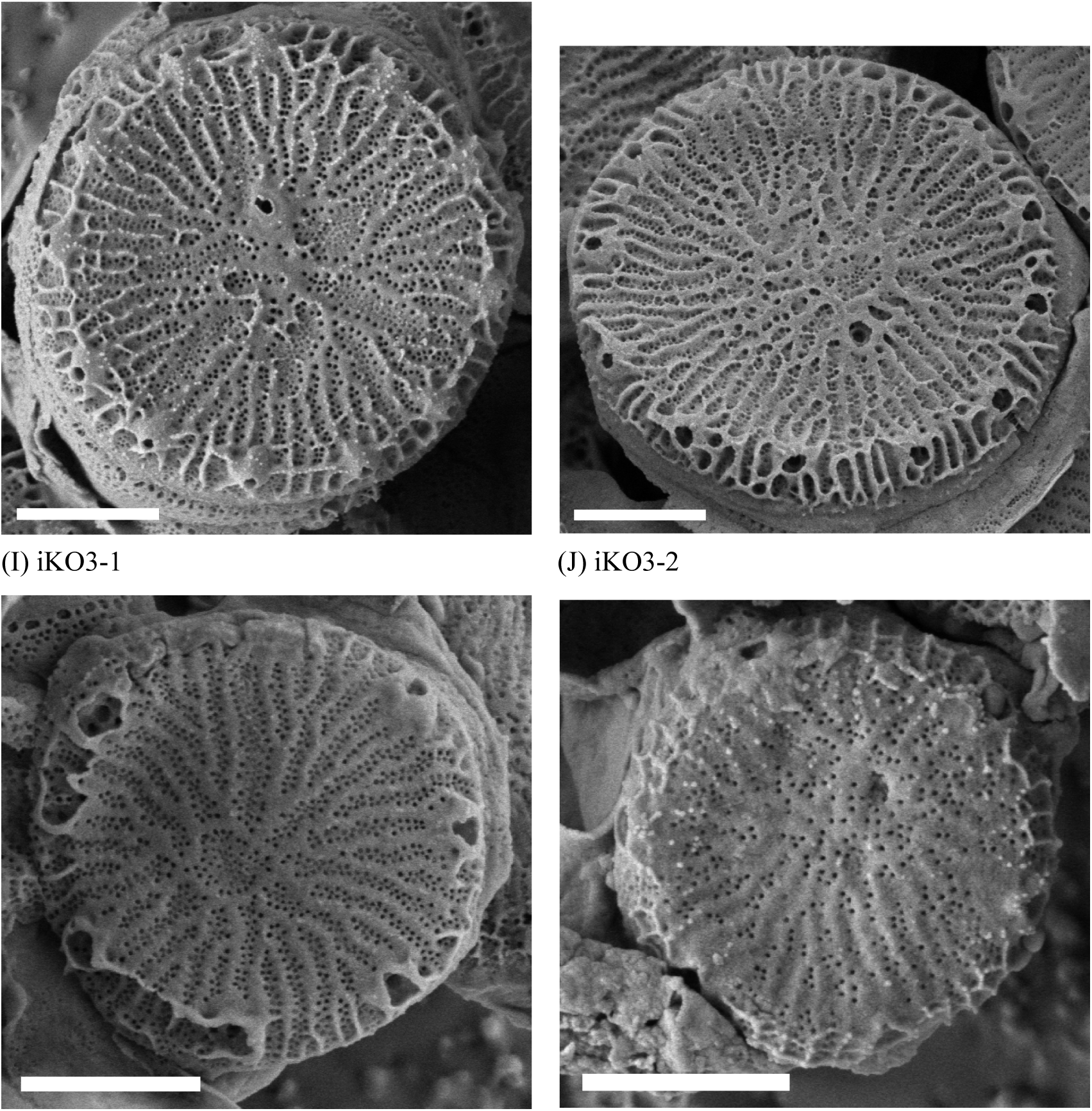
Representative scanning electron microscopy (SEM) images of frustule valves from wild-type and genetically transformed *T. pseudonana* strains. (A) OE1-1 and (B) OE1-2: *TpSil1* overexpression strains. (C) KO1/2-1 and (D) KO1/2-2: *TpSil1* and *TpSil2* double*-*knockout strains. (E) and (F) WT: wild-type strain. (G) OE3-1 and (H) OE3-2: *TpSil3* overexpression strains, (I) iKO3-1 and (J) iKO3-2: incomplete *TpSil3* knockout strains. Key structural elements are marked as follows: Costae: green lines; Cross-connections: orange lines; Areola pores: red rectangles; Cribrum pores: blue dots; Fultoportulae: yellow circles; Satellite pores of fultoportula: yellow arrows; Central pore of fultoportula: light green arrow. Scale bar: 1 μm.

To rigorously characterize morphological adaptations induced by genetic modifications on *silaffins*, structural features were quantified by analyzing 12 individual frustules per strain. The WT frustules had an average valve diameter of 3.75 ± 0.35 μm (Fig. 4A), while iKO3 frustules exhibited a significant reduction (*P* < 0.001) by 19.33–23.94%. Marginal fultoportulae counts (Fig. 4B) were reduced in KO1/2 (*P* < 0.01) and iKO3 (*P* < 0.001), with iKO3 showing the most significant reduction (*P* < 0.001) in central fultoportulae counts (Fig. 4C). The mean fultoportula area (Fig. 4D) did not significantly change in OE1, OE3, and iKO3, but KO1/2 exhibited a significant increase compared to WT (*P* < 0.001), suggesting an altered silica distribution around the fultoportulae, consisting with the localization of TpSil1-GFP (Poulsen et al., 2013). Cribrum pore density (Fig. 4E) was significantly higher in KO1/2 (*P* < 0.05), but reduced in iKO3 (*P* < 0.01). Pore area (Fig. 4F) was smaller in OE3, KO1/2, and iKO3 compared to WT (*P* < 0.001), highlighting the role of *TpSil1/2* and *TpSil3* in modulating pore size. The pore-to-total-area ratio (Fig. 4G) was reduced in both OE3 and iKO3 compared to WT (*P* < 0.001), further indicating that *TpSil3* affects silica distribution.

**Fig. 4.**
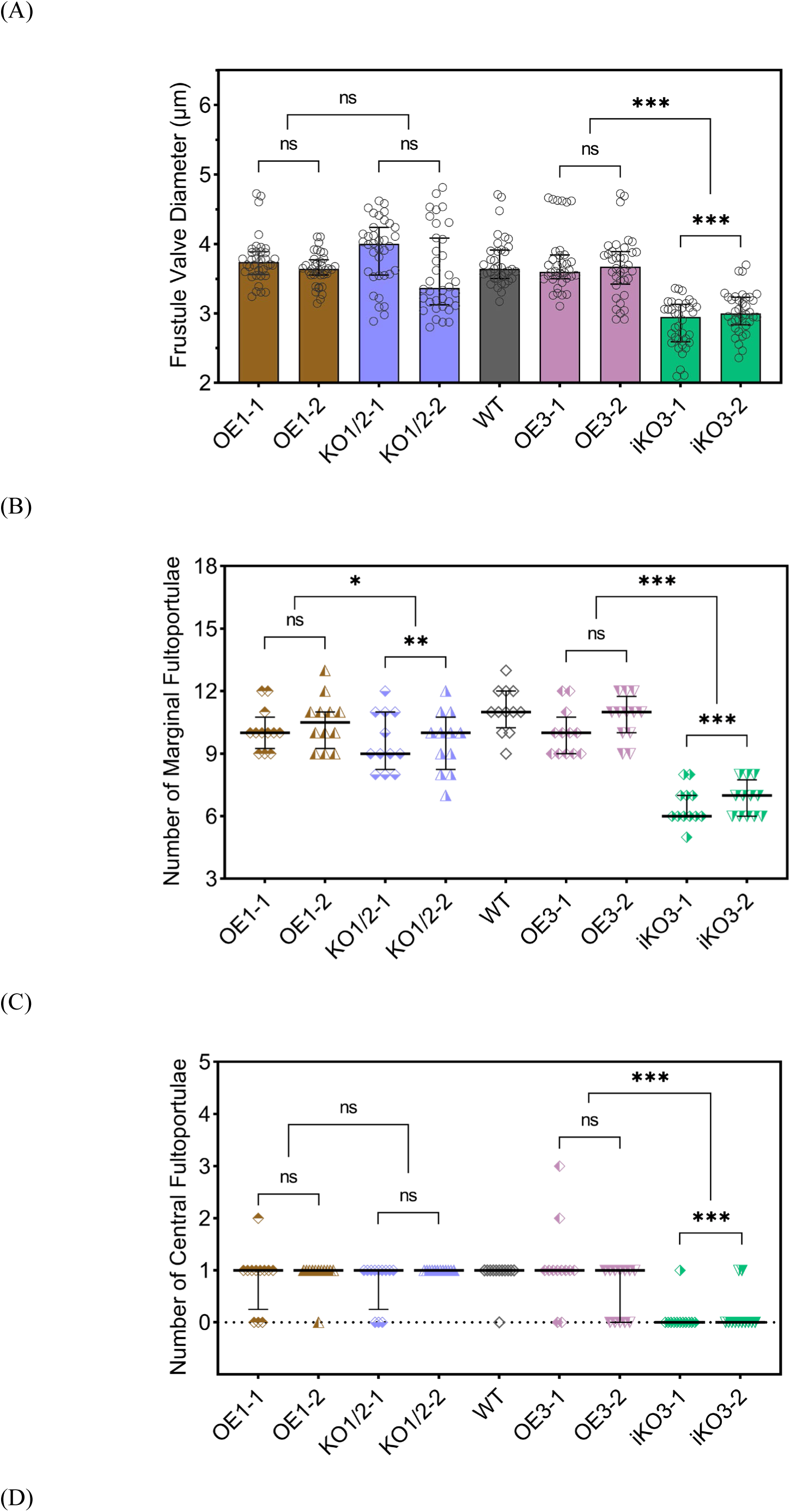
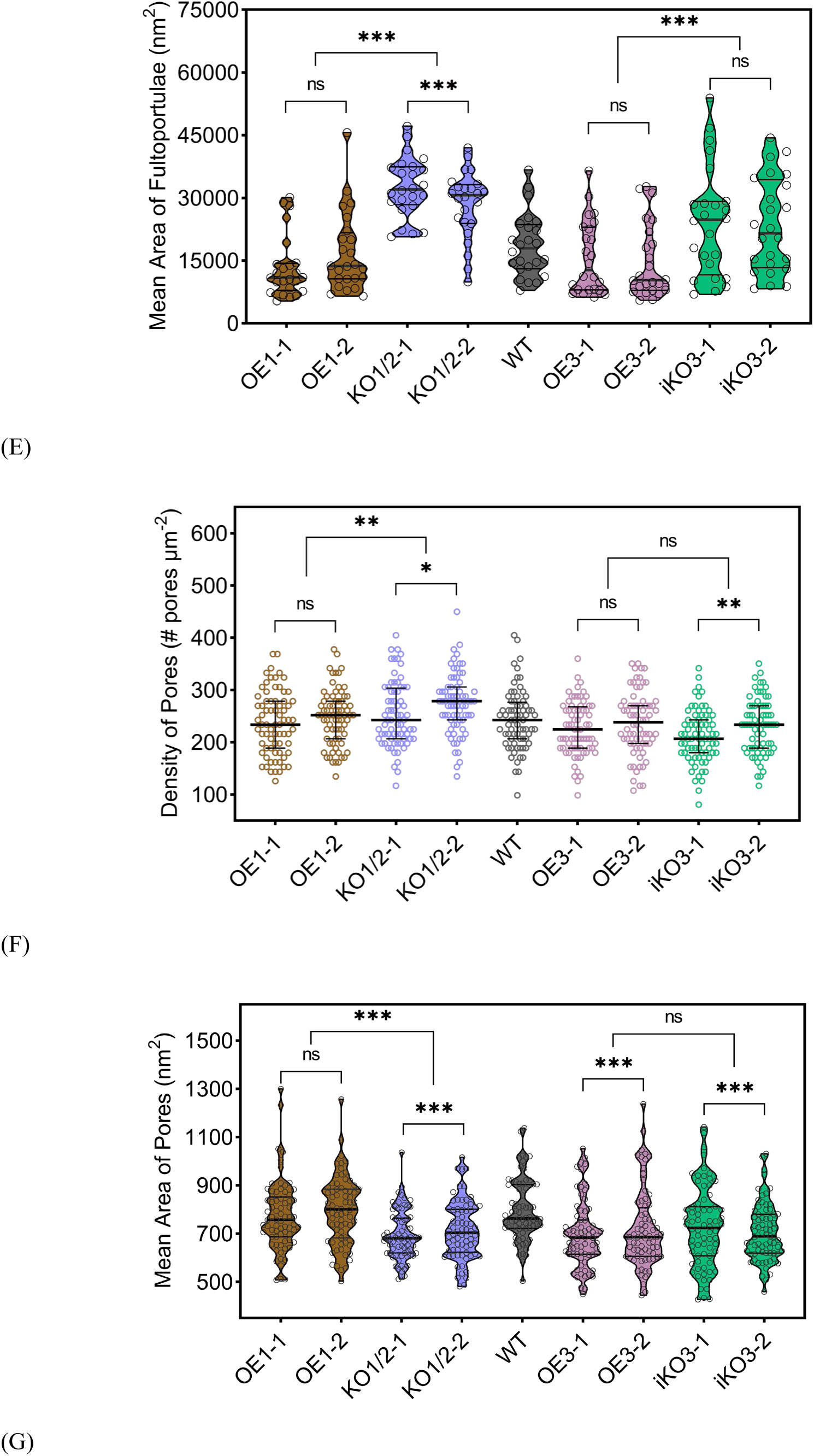
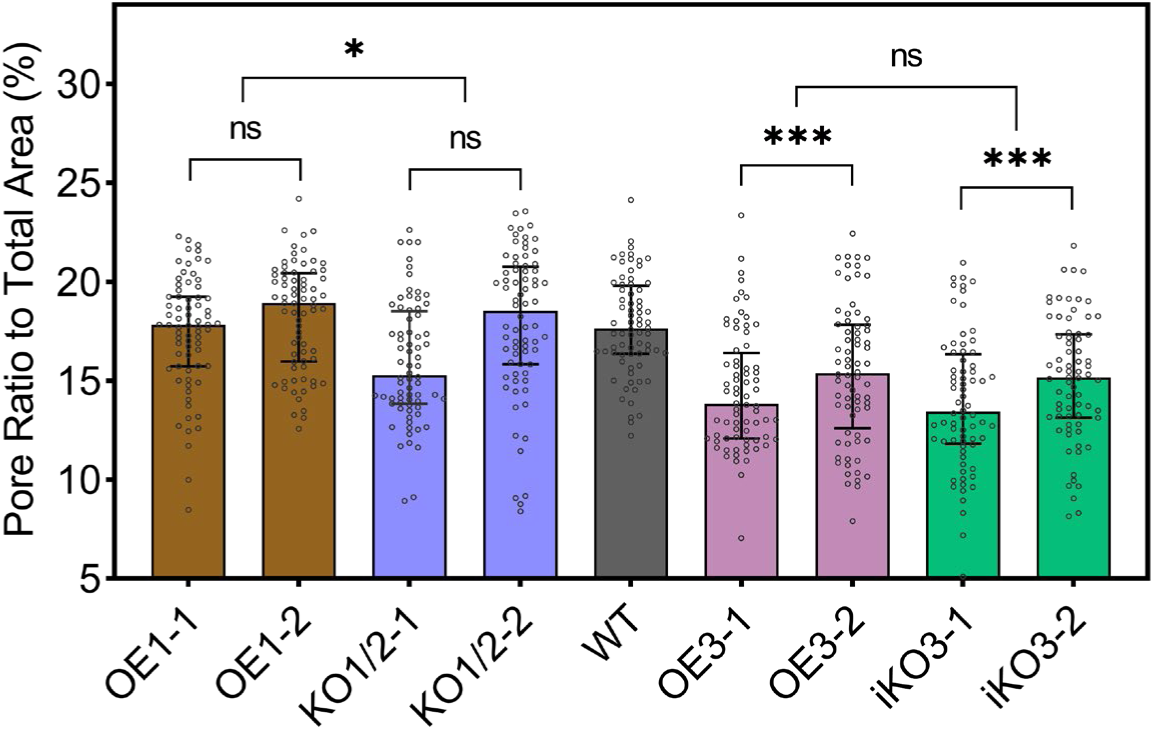
Quantitative analysis on morphological characteristics of frustule valves from wild-type and genetically transformed *T. pseudonana* strains. (A) Frustule valve diameter, calculated as the mean of 3 measurements per frustule, across 12 frustules per strain (*n* = 36). (B) Number of marginal fultoportulae per frustule (*n* = 12 frustules). (C) Number of central fultoportulae per frustule (*n* = 12 frustules). (D) Mean fultoportula area, calculated from two front-facing fultoportulae per frustule, across 12 frustules per strain (*n* = 24). (E) Cribrum pore density, (F) Mean cribrum pore area, and (G) Cribrum pore-to-total-area ratio, calculated from six 60 × 60-pixel regions per frustule SEM image, across 12 frustules per strain (*n* = 72). Strains: WT: wild-type strain; OE1-1 and OE1-2: *TpSil1* overexpression strains; KO1/2-1 and KO1/2-2: *TpSil1* and *TpSil2* double-knockout strains; OE3-1 and OE3-2: *TpSil3* overexpression strains; iKO3-1 and iKO3-2: incomplete *TpSil3* knockout strains. Thin lines represent the quartiles, the thick line marks the median, and individual data points are displayed as dots within the plots. Statistical comparisons were conducted between two groups using an independent two-sample t-test with equal variance assumptions (\**P* < 0.05, \*\**P* < 0.01, \*\*\**P* < 0.001).

In summary, overexpression of *silaffins* had little impact on valve surface characteristics, except in OE3 with reduced cribrum pore area. In KO1/2 strains, the increased fultoportula area, reduced marginal fultoportulae number, enhanced cribrum pore density and smaller cribrum pore areas suggest that *TpSil1/2* are involve in fultoportulae morphogenesis and fine-tune cribrum pore distribution. In contrast, *TpSil3* disruption resulted in remarkable reductions in valve diameter, fultoportulae number, cribrum pore densities and areas, indicating that *TpSil3* plays a broader role in regulating both global frustule scale and local morphology.

### 3.4 Silaffins on Physicochemical and Optical Properties of Bulk Frustule

The morphological analysis of individual frustule reveals invaluable insights into silica deposition and patterning. However, the properties of bulk frustules are influenced not only by individual frustules but also by the interactions and spatial arrangements within larger assemblies. This section examines how genetic modifications on *silaffins* affect the physicochemical and optical properties of bulk frustules.

BET surface area analysis (Table 1) revealed that KO1/2 frustules had the highest surface area, followed by OE1 and OE3, which were all significantly higher than WT. iKO3 frustules displayed a slight, statistically insignificant increase in surface area. Similarly, KO1/2 and OE1 exhibited significantly higher pore volumes compared to WT, while OE3 and iKO3 frustules had lower pore volumes, but still exceeded WT (Table 1). OE1 showed the largest average pore diameter, followed by KO1/2, both larger than WT. In contrast, OE3 and iKO3 had smaller diameters. Laser diffraction particle size analysis in ethanol (Fig. 5A) revealed that OE1 exhibited a narrower particle size range (4.54–17.60 µm, indicated by Dv(10) and Dv(90)) compared to WT, while OE3 displayed the smallest size distribution (1.73–7.12 µm). In contrast, KO1 showed broader distribution (5.39–48.90 µm), and iKO3 had the widest distribution (10.90–62.70 µm). These results indicate that overexpression of *silaffins* promotes finer, more homogeneous dispersion, while knockout lead to aggregation and a wider size distribution.

**Fig. 5.**
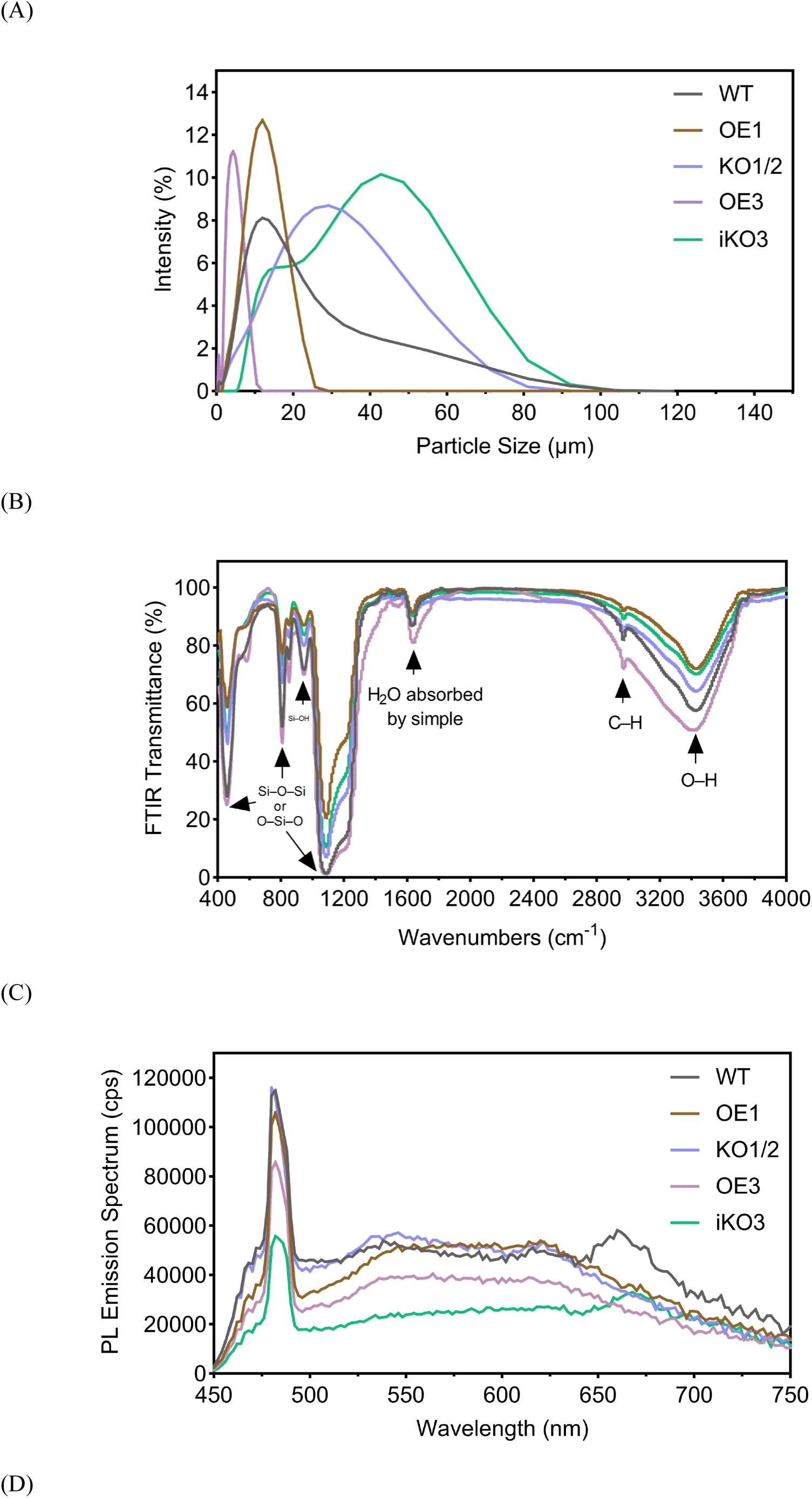
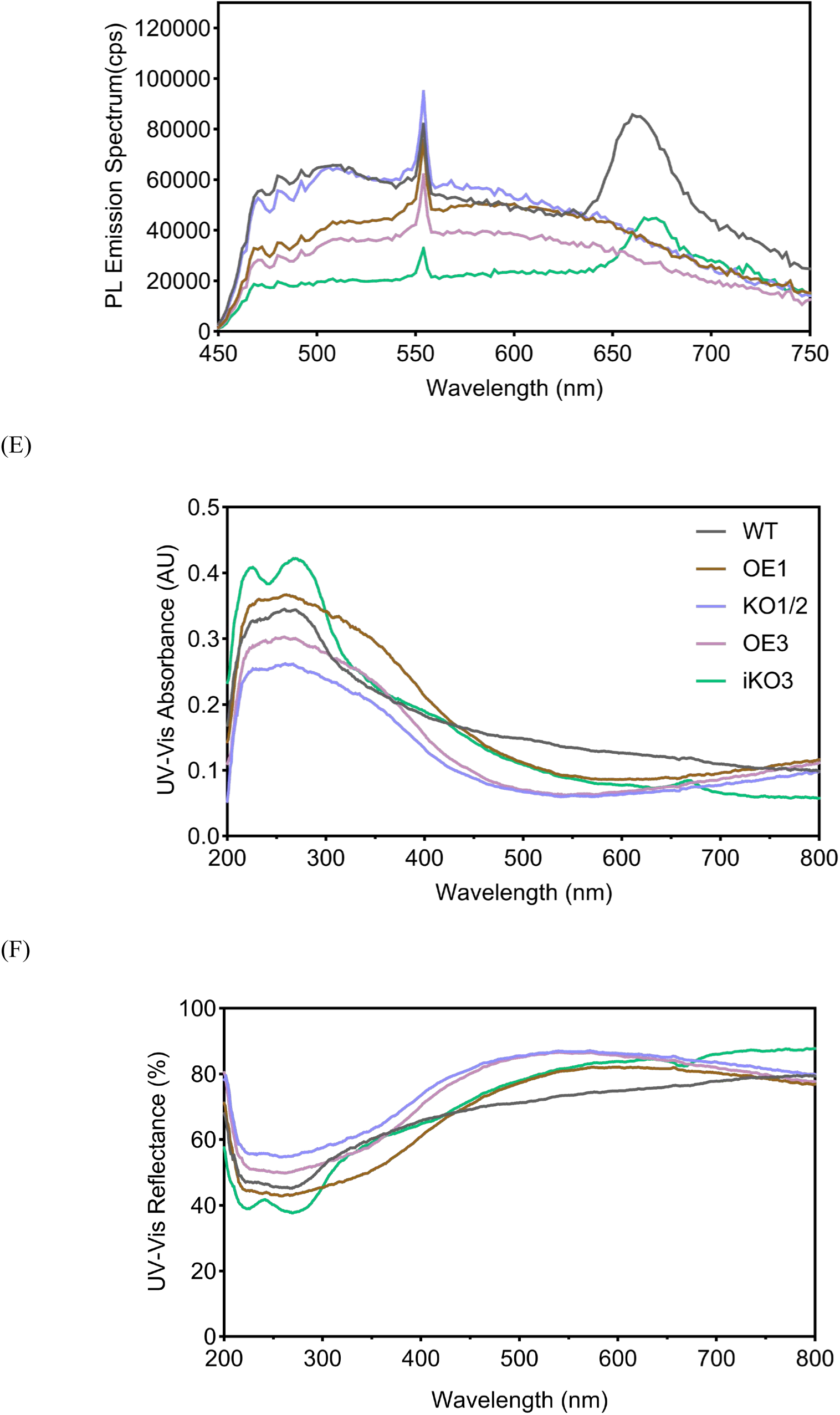
Physicochemical and optical properties of *T. pseudonana* frustules for wild-type and genetically transformed strains. (A) Laser diffraction particle size distribution. (B) Fourier transform infrared (FTIR) spectroscopy analysis. (C) Photoluminescence (PL) emission spectra under 330 nm excitation, measured in counts per second (cps). (D) PL emission spectra under 380 nm excitation, measured in cps. (E) UV-visible diffuse reflectance spectroscopy (UV-Vis DRS) absorbance spectra, recorded in absorbance units (AU). (F) UV-Vis DRS reflectance spectra. Each measurement represents the average of three biologically independent samples. Strains: WT: wild-type strain; OE1: OE1-1, *TpSil1* overexpression strain; KO1/2: KO1/2-1, *TpSil1* and *TpSil2* double-knockout strain; OE3: OE3-1, *TpSil3* overexpression strain; iKO3: iKO3-1, incomplete *TpSil3* knockout strain.

**Table 1.**
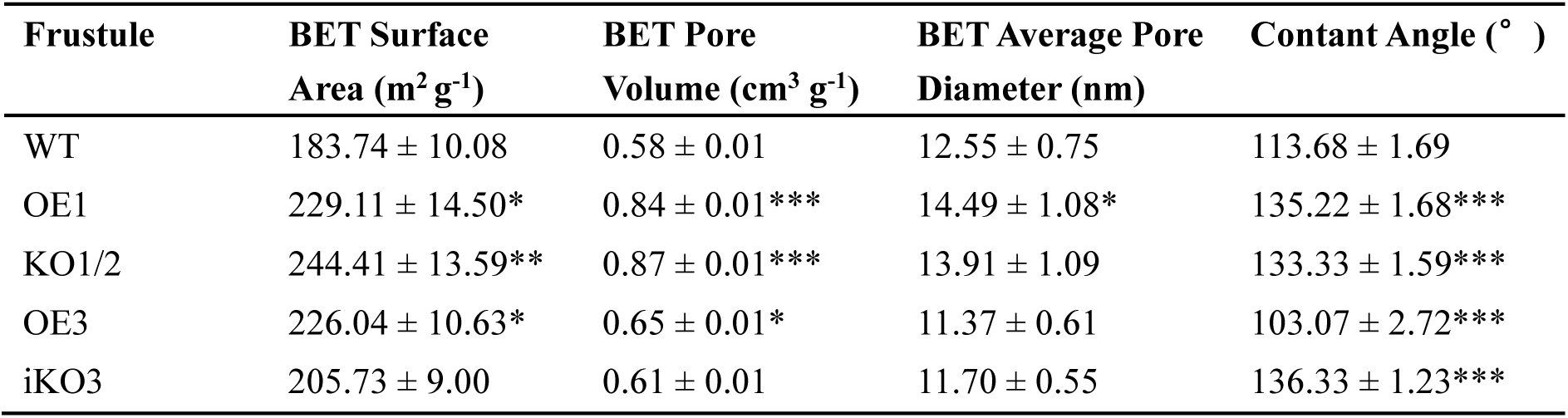
Brunauer-emmett-teller (BET) and contact angle analyses of bulk frustules for wild-type and genetically transformed *T. pseudonana* strains. Strains: WT: wild-type strain; OE1: OE1-1, *TpSil1* overexpression strain; KO1/2: KO1/2-1, *TpSil1* and *TpSil2* double-knockout strain; OE3: OE3-1, *TpSil3* overexpression strain; iKO3: iKO3-1, incomplete *TpSil3* knockout strains. Data are presented as mean ± standard deviation (SD), with n = 3 biologically independent samples. Statistical comparisons were performed against the wild-type strain using an independent two-sample t-test with equal variance assumptions (\**P* < 0.05, \*\**P* < 0.01, \*\*\**P* < 0.001).

| Frustule | BET Surface Area (m<sup>2</sup> g<sup>-1</sup>) | BET Pore Volume (cm<sup>3</sup> g<sup>-1</sup>) | BET Average Pore Diameter (nm) | Contact Angle (°) |
| --- | --- | --- | --- | --- |
| WT | 183.74 ± 10.08 | 0.58 ± 0.01 | 12.55 ± 0.75 | 113.68 ± 1.69 |
| OE1 | 229.11 ± 14.50* | 0.84 ± 0.01*** | 14.49 ± 1.08* | 135.22 ± 1.68*** |
| KO1/2 | 244.41 ± 13.59** | 0.87 ± 0.01*** | 13.91 ± 1.09 | 133.33 ± 1.59*** |
| OE3 | 226.04 ± 10.63* | 0.65 ± 0.01* | 11.37 ± 0.61 | 103.07 ± 2.72*** |
| iKO3 | 205.73 ± 9.00 | 0.61 ± 0.01 | 11.70 ± 0.55 | 136.33 ± 1.23*** |

Fourier transform infrared spectroscopy (FTIR) (Fig. 5B) confirmed that silicon is the primary structural component of frustules, with characteristic peaks for Si–O–Si and O–Si–O vibrations at 460, 800, and 1100 cm^-1^. While no novel peaks were observed in the genetically modified frustules, variations in peak intensities indicated changes in surface chemistry. OE3 and WT frustules exhibited lower transmittance, suggesting more intricate silica networks with higher infrared-absorbing functional groups. The frustules of iKO3, OE1 and KO1/2 exhibited significantly higher contact angles compared to WT (Table 1), indicating increased surface hydrophobicity. Conversely, OE3 showed a significantly lower contact angle, suggesting enhanced surface hydrophilicity. The observed changes in surface wettability are likely attributable to modifications in surface chemistry, as evidenced by the FTIR analysis (Fig. 5B). The peak at 3400 cm^-1^ corresponds to O–H stretching vibrations, and the increase in O–H groups can enhance surface hydrophilicity.

Photoluminescence (PL) spectra (Fig. 5C, D) showed that WT frustules exhibited the highest PL intensity under both 330 nm and 380 nm excitation wavelengths, suggesting a high density of emissive centers and an optimal structure for luminescence. KO1/2 frustules also showed strong PL under 380 nm excitation. In contrast, OE1, OE3, and iKO3 frustules displayed reduced PL intensities, potentially attributable to increased non-radiative losses in their structures. Both WT and iKO3 frustules exhibited red emission tails (650–700 nm), possibly related to defect emissions or energy traps (Choudhury and Choudhury, 2013). UV-visible diffuse reflectance spectroscopy (UV-Vis DRS) (Fig. 5E, F) revealed that all frustules strongly absorbed UV light (220–380 nm) while transmitting visible light. iKO3 and OE1 exhibited higher UV absorbance and visible light reflectance compared to WT, suggesting enhanced UV protection and increase visible light scattering. Conversely, KO1/2 and OE3 demonstrated higher reflectance than WT across most of the UV-Vis range, with a peak reflectance of over 87% around 550 nm, indicating structures optimized for light scattering. In summary, these analyses demonstrate how genetic engineering, through the overexpression or knockout of *silaffins*, influences the physicochemical and optical properties of frustules, highlighting the potential for diverse applications, including catalysis, adsorption, UV protection, and light scattering.

## 4. Discussions

This study explored the roles of two key *silaffin* genes of *T. pseudonana*, *TpSil1* and *TpSil3*, in regulating diatom physiology, frustule morphology and bulk frustule properties through gene overexpression and CRISPR/Cas9-mediated knockout approaches. Overexpression was achieved with the *NR* promoter, leading to 2–4 fold increased expression of *TpSil1* (Fig. 1B), or 16–22 fold increased expression of *TpSil3* (Fig. 1C). The discrepancy in expression levels may arise from factors such as the lower baseline expression of *TpSil3*, feedback regulation limiting *TpSil1* upregulation, or chromosomal positioning effects on gene expression (Becskei et al., 2005). Given the concentration-dependent silica deposition activity of TpSil1/2H and TpSil3 *in vitro* (Poulsen and Kröger, 2004), varying expression levels of *TpSil1/2* or *TpSil3* genes may exert distinct effects on diatom silicification, which require further investigation using different expression cassettes and site-specific gene expression.

CRISPR/Cas9-mediated knockout of *TpSil1* and *TpSil3* was performed using dual sgRNAs. Notably, one sgRNA (*TpSil1*_sgRNA1, SI Appendix, Fig. S4 and Table S3) targeting *TpSil1* also targeted *TpSil2* due to their high sequence homology, resulting in homozygous knockout strains in which both genes were disrupted. However, generating homozygous *TpSil3* knockout strains proved challenging. The reduced yield of sub-clones during re-plating, incomplete gene disruption observed in 168 analyzed sub-clones (SI Appendix, Supplementary Text S2.2.2), and the broad subcellular localization of *TpSil3* across the frustule surface (Poulsen et al., 2013) collectively indicate that the homozygous knockout of *TpSil3* is lethal, underscoring the essential, non-redundant role of *TpSil3* in *T. pseudonana*.

The physiological role of frustules in diatoms remains speculative, with proposed functions including UV protection (Aguirre et al., 2018), light scattering (Goessling et al., 2018), and nutrient uptake (Mitchell et al., 2013). In this study, we explored the connection between frustule morphology and diatom physiological processes through genetic manipulation of frustule-related genes, *silaffins*. i) Growth and photosynthesis: disruptions of *TpSil1/2* or *TpSil3* resulted in significantly increased growth rate (Fig. 2A) and enhanced photosynthetic activity (Fig. 2B, C, D), coupled with decreased cellular silica content (Fig. 2E). Reduced silicon content and increased growth rates have also been reported in *TpSin1* knockout (Görlich et al., 2019) and *CcSAP2* knockdown strains (Wang et al., 2023), suggesting that decreased silica deposition may reduce energy cost, allowing metabolic resources to be redirected towards growth. ii) Nutrient uptake: the reduced cell size observed in iKO3 strains (Fig. 2F, 3I, J, and 4A) could increase the surface area-to-volume ratio, potentially facilitating improved nutrient uptake efficiency and supporting accelerated growth rates. iii) Altered optical properties: iKO3 frustules exhibited increased UV shielding and greater visible light transmittance (Fig. 5E, F), which may improve light utilization and further boost photosynthetic efficiency. Together, these data demonstrate that genetic manipulation of frustule-related genes can significantly influence diatom metabolism, providing a basis for technological strategies to optimize metabolic processes in diatoms through modulation of frustule structures.

Quantitative analysis of frustule valve morphology revealed the distinct yet complementary roles of *TpSil1/2* and *TpSil3* in frustule valve morphogenesis. The morphological changes in frustules induced by genetic modifications to date are summarized in Table 2. In the microscale, disruption of *TpSil3* resulted in significant reductions in overall cell size (Fig. 2F) and valve diameter (Fig. 4A). Similar reductions were observed in the knockdown of *CcSAP2* in *C. cryptica* (Wang et al., 2023). Conversely, knockout or knockdown of *TpSilacidin* in *T. pseudonana* displayed significant increases in valve size (Kirkham et al., 2017; Belshaw et al., 2023). These findings suggest that TpSil3 and CcSAP2 act in a concert way to promote valve development, enabling the frustule to reach its full size, while *TpSilacidin* serves as a negative regulator, restricting valve expansion.

**Table 2.**
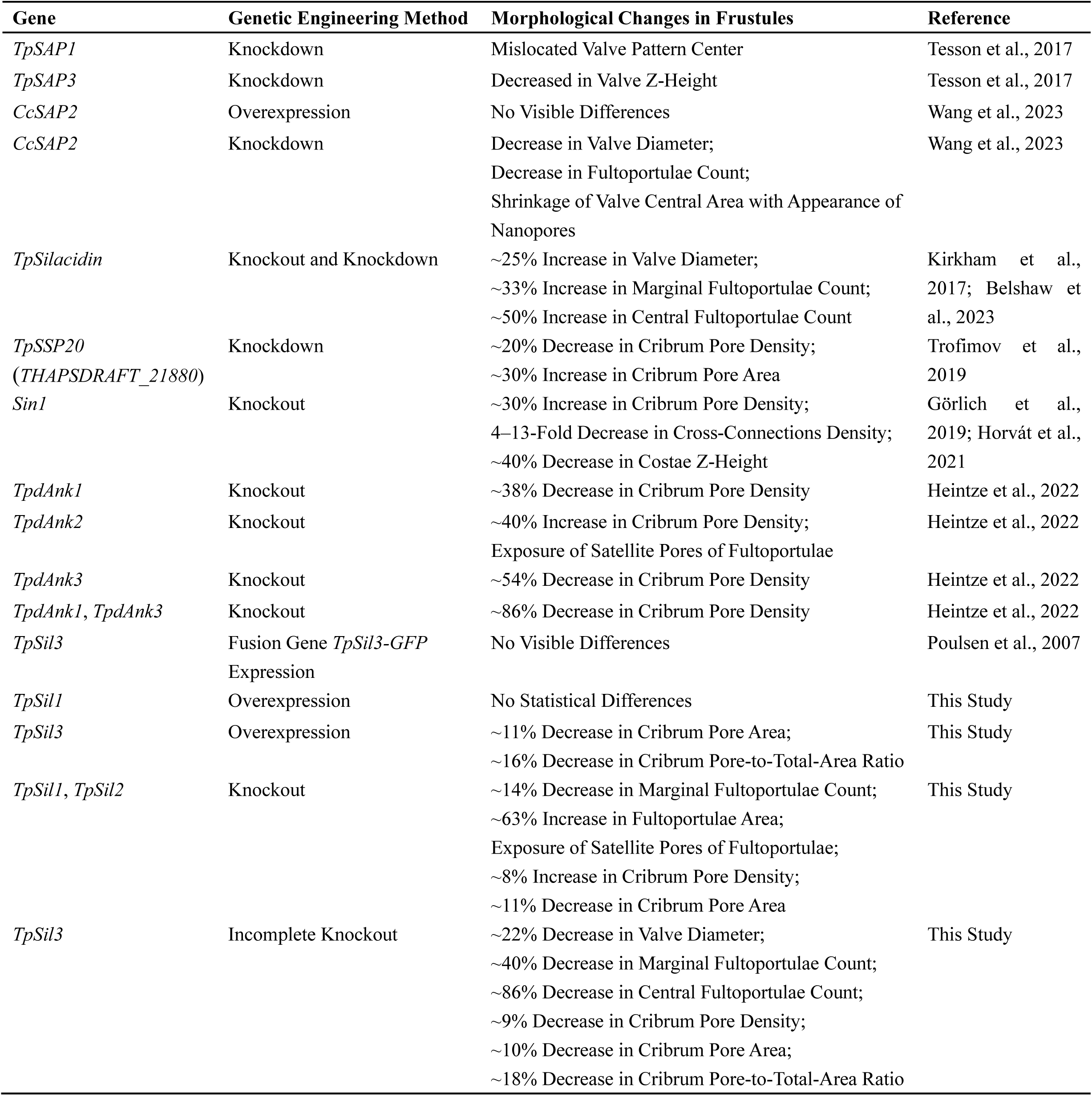
Genetic engineering-induced changes in morphology of diatom frustules. Comparisons are made between frustules of genetically modified and wild-type diatom strains. Cc, *Cyclotella cryptica*; Tp, *Thalassiosira pseudonana*; SAP, silicalemma associated protein; SSP, soluble silicome protein; Sin1, Silicanin-1; dAnk, diatom ankyrin; Sil, silaffin; GFP, green fluorescent protein.

| Gene | Genetic Engineering Method | Morphological Changes in Frustules | Reference |
| --- | --- | --- | --- |
| *TpSAP1* | Knockdown | Mislocated Valve Pattern Center | Tesson et al., 2017 |
| *TpSAP3* | Knockdown | Decreased in Valve Z-Height | Tesson et al., 2017 |
| *CcSAP2* | Overexpression | No Visible Differences | Wang et al., 2023 |
| *CcSAP2* | Knockdown | Decrease in Valve Diameter;<br>Decrease in Fultoportulae Count;<br>Shrinkage of Valve Central Area with Appearance of Nanopores | Wang et al., 2023 |
| *TpSilacidin* | Knockout and Knockdown | ~25% Increase in Valve Diameter;<br>~33% Increase in Marginal Fultoportulae Count;<br>~50% Increase in Central Fultoportulae Count | Kirkham et al., 2017; Belshaw et al., 2023 |
| *TpSSP20*<br>(*THAPSDRAFT_21880*) | Knockdown | ~20% Decrease in Cribrum Pore Density;<br>~30% Increase in Cribrum Pore Area | Trofimov et al., 2019 |
| *Sin1* | Knockout | ~30% Increase in Cribrum Pore Density;<br>4–13-Fold Decrease in Cross-Connections Density;<br>~40% Decrease in Costae Z-Height | Görlich et al., 2019; Horvát et al., 2021 |
| *TpdAnk1* | Knockout | ~38% Decrease in Cribrum Pore Density | Heintze et al., 2022 |
| *TpdAnk2* | Knockout | ~40% Increase in Cribrum Pore Density;<br>Exposure of Satellite Pores of Fultoportulae | Heintze et al., 2022 |
| *TpdAnk3* | Knockout | ~54% Decrease in Cribrum Pore Density | Heintze et al., 2022 |
| *TpdAnk1, TpdAnk3* | Knockout | ~86% Decrease in Cribrum Pore Density | Heintze et al., 2022 |
| *TpSil3* | Fusion Gene *TpSil3-GFP*<br>Expression | No Visible Differences | Poulsen et al., 2007 |
| *TpSil1* | Overexpression | No Statistical Differences | This Study |
| *TpSil3* | Overexpression | ~11% Decrease in Cribrum Pore Area;<br>~16% Decrease in Cribrum Pore-to-Total-Area Ratio | This Study |
| *TpSil1, TpSil2* | Knockout | ~14% Decrease in Marginal Fultoportulae Count;<br>~63% Increase in Fultoportulae Area;<br>Exposure of Satellite Pores of Fultoportulae;<br>~8% Increase in Cribrum Pore Density;<br>~11% Decrease in Cribrum Pore Area | This Study |
| *TpSil3* | Incomplete Knockout | ~22% Decrease in Valve Diameter;<br>~40% Decrease in Marginal Fultoportulae Count;<br>~86% Decrease in Central Fultoportulae Count;<br>~9% Decrease in Cribrum Pore Density;<br>~10% Decrease in Cribrum Pore Area;<br>~18% Decrease in Cribrum Pore-to-Total-Area Ratio | This Study |

To the mesoscale, fultoportulae are hollow cylindrical channels spanning from the proximal to distal valve surfaces, facilitating the secretion of β-chitin (Herth, 1979). The number of fultoportulae is significantly reduced in *TpSil3* incomplete knockout and *CcSAP2* knockdown frustules (Wang et al., 2023), whereas it significantly increases in *TpSilacidin* knockout or knockdown frustules (Kirkham et al., 2017; Belshaw et al., 2023), suggesting a correlation between valve size and fultoportulae number. Additionally, in WT frustules, fultoportulae are typically covered by funnel-shaped tubes. However, in KO1/2 frustules, these tubes are absent, leaving the satellite pores exposed (Fig. 3C, D). Similar observations have been reported in *TpdAnk2* knockout frustules (Heintze et al., 2022). Furthermore, GFP-tagged TpSSP11 (THAPSDRAFT_5357) (Fattorini and Maier, 2022), TpSSP3 (THAPSDRAFT_22220), TpSSP16 (THAPSDRAFT_8219) and TpSSP20 (THAPSDRAFT_21880) (Skeffington et al., 2022) exhibit characteristic localization patterns at the fultoportulae, indicating the potential involvement of these proteins in fultoportulae morphogenesis. These findings highlight the complexity and hierarchy of the regulatory network governing fultoportula development. Disruption of a single gene within this network can result in significant alterations to frustule morphology.

Regarding the cribrum pore pattern morphogenesis, knockout of *TpSin1* (Görlich et al., 2019; Horvát et al., 2021) or *TpdAnk2* (Heintze et al., 2022) resulted in ∼35% or ∼40% increases in valve pore density, respectively. In contrast, knockout of *TpdAnk1* or *TpdAnk3* caused significant decreases of ∼38% or ∼54% (Heintze et al., 2022). In this study, knockout of *TpSil1/2* caused a modest 8% increase in cribrum pore density, while disruption of *TpSil3* led to a slight 9% decrease. These results suggest that silaffins may not directly regulate cribrum pore morphogenesis, but instead influence it indirectly through interactions with other regulatory proteins, such as TpdAnks and TpSin1. TpdAnks localized on the exterior of SDV membrane (Heintze et al., 2022), TpSin1 spans the SDV membrane (Kotzsch et al., 2017), and silaffins are part of the organic matrix inside the SDV lumen (Poulsen et al., 2013). Given their distinct spatial locations, these proteins likely operate within a coordinated regulatory network to govern pore patterning. How these proteins interact to establish pore patterns remains an open question, highlighting a sophisticated interplay that may involve LCPAs.

TpSil3 isolated from frustule silica matrix forms flat sheet-like silica structures *in vitro* with LCPAs (Poulsen and Kröger, 2004), and its transcriptional pattern is specifically correlated with valve synthesis (Frigeri et al., 2006), suggesting its role in base layer formation. The lethal phenotype in homozygous knockouts and the micro- and mesoscale morphological changes in the incomplete knockouts underscore the critical role of TpSil3 in coordinating frustule morphogenesis. TpSil1/2H generate spherical silica structures *in vitro* with LCPAs (Poulsen and Kröger, 2004), and *TpSil1* transcription is induced during girdle band formation (Frigeri et al., 2006). Structural changes in the fultoportulae and cribrum pore density of homozygous knockouts highlight the role of TpSil1/2 in modulating specific mesoscale features of the frustule. Together, these results underline the complementary yet distinct roles of *TpSil1/2* and *TpSil3* in orchestrating the complex process of frustule morphogenesis at multiple scales.

For the overexpression strains, except for the *TpSil3* overexpression strains that exhibited a reduction in cribrum pore area, no significant visible or quantitative changes were observed in frustule morphology. However, functional alterations in bulk frustules of overexpression strains were evident, indicating that overexpression may influence the girdle band morphology, internal frustule structures, or the spatial interactions between girdle bands and valves, ultimately impacting overall frustules performance. Table 3 summarizes the currently known genetic engineering-induced modifications to frustule functionality and performance.

**Table 3.** Genetic engineering-induced changes in functionality or performance of diatom frustules. Comparisons are made between frustules of genetically modified and wild-type diatom strains. Cc, *Cyclotella cryptica*; Tp, *Thalassiosira pseudonana*; Sin1, Silicanin-1; Sil, silaffin; SAP, silicalemma associated protein; YFP, yellow fluorescent protein; GOx, glucose oxidase; PL, photoluminescence.

| Gene | Genetic Engineering Method | Modified Frustule Properties | Reference |
| --- | --- | --- | --- |
| *TpSin1* | Knockout | Compromised Strength and Stiffness | Görlich et al., 2019 |
| *TpSil3, Sin1* | Fusion Gene<br>*TpSil3-YFP-GOx*<br>Expression in *TpSin1*<br>Knockout Strain | Valve Surface Exhibits GOx Catalytic Activity;<br>~56% Increase in GOx Catalytic Activity in *TpSin1* Knockout Frustule | Kumari et al., 2020 |
| *CcSAP2* | Knockdown | ~51% Increase in BET Surface Area;<br>~36% Increase in BET Pore Volume;<br>~15% Increase in Liquid Absorption Ratio;<br>Enhanced Coagulation and Hemostatic Effects | Wang et al., 2023 |
| *TpSil1* | Overexpression | ~24% Increase in BET Surface Area;<br>~45% Increase in BET Pore Volume;<br>More Hydrophobic Surface;<br>Smaller and More Uniform Particle Size Distribution in Liquid;<br>Reduced PL Intensity under 380 nm Excitation;<br>Higher UV Absorbance and Visible Light Reflectance | This Study |
| *TpSil3* | Overexpression | ~23% Increase in BET Surface Area;<br>~12% Increase in BET Pore Volume;<br>More Hydrophilic Surface;<br>Smaller and More Uniform Particle Size Distribution in Liquid;<br>Reduced PL Intensity under 330 and 380 nm Excitation;<br>Higher UV-Visible Light Reflectance | This Study |
| *TpSil1* and *TpSil2* | Knockout | ~32% Increase in BET Surface Area;<br>~50% Increase in BET Pore Volume;<br>More Hydrophobic Surface;<br>Larger and More Aggregated Particulate Dispersion in Liquid;<br>Slightly Enhanced PL Intensity under 380 nm Excitation;<br>Higher UV-Visible Light Reflectance | This Study |
| *TpSil3* | Incomplete Knockout | ~50% Increase in BET Pore Volume;<br>More Hydrophobic Surface;<br>Larger and More Aggregated Particulate Dispersion in Liquid;<br>Higher UV Absorbance and Visible Light Reflectance;<br>Reduced PL Intensity under 330 and 380 nm Excitation | This Study |

Knockout of *TpSin1* compromised the strength and stiffness of frustules, possibly due to the decreased cross-connection density on the valve and the reduced silica deposition (Görlich et al., 2019). Similarly, disruptions of *TpSil1/2* or *TpSil3* led to absence of cross-connection features (Fig. 3) and lower cellular silica content (Fig. 2E), suggesting diminished mechanical stability in KO1/2 and iKO3 frustules. Furthermore, the expression of the fusion gene *TpSil3-YFP-GOx* in *TpSin1* knockout strains enhanced the catalytic activity of glucose oxidase (GOx) immobilized on valves by ∼55% compared to WT frustules (Kumari et al., 2020). Given the similarities in frustule morphologies (e.g., increased cribrum pore density, reduced cross-connection density, unchanged valve size) between *TpSil1/2* and *TpSin1* knockouts, KO1/2 frustules likely exhibit enhanced catalytic capability of enzyme immobilized on the valve surface.

BET surface area and liquid absorbability have been identified as key factors influencing the hemostatic performance of frustules (Wang et al., 2023). Frustules of OE3 are likely to demonstrate superior hemostatic effects compared to WT, due to the increased BET surface area and enhanced surface hydrophobicity (Table 1). Additionally, *TpSil1* and *TpSil3* overexpression frustules exhibited smaller and more homogeneous dispersion in liquid, potentially improving their efficiency in applications such as catalysis, adsorption, and drug delivery.

Genetic modifications of *silaffins* also affected the optical properties of frustules. Notably, OE1 and iKO3 frustules exhibited elevated UV absorbance and visible light reflectance (Fig. 5E, F), making them well-suited for applications requiring efficient UV absorption and visible light transmission, such as UV-blocking coatings. In contrast, OE3 and KO1/2 frustules, with enhanced UV and visible light reflectance, could be ideal candidates for light-reflective applications. WT frustules, exhibited the strongest light-emitting properties among tested frustules (Fig. 5C, D), hold promise for use as optically active materials. Collectively, genetic modifications of *silaffins* significantly altered the bulk properties of frustule, potentially enhancing their hemostatic, catalytic, optical, and light-reflective performances, thereby making them versatile candidates for a wide range of biotechnological and industrial applications.

In summary, this study elucidates the roles of two representative *silaffin* genes (*TpSil1* and *TpSil3*) in regulating diatom physiology, individual frustule morphology and bulk frustule functionality. Future omics analysis, including proteomics and transcriptomics of silaffin mutant strains, are anticipated to uncover additional proteins involved in frustule morphogenesis. Additionally, assessing the impact of *silaffin* mutations on other valuable diatom metabolites, such as fucoxanthin, chitin, and polyunsaturated fatty acids, could deepen our understanding of frustule contributions to diatom physiology and maximize the biotechnological potential of these engineered strains. Lastly, large-scale diatom cultivation and industrial-scale testing of frustule properties will be essential to evaluate their practical applications and commercial viability.

## 5. Conclusion

This study investigated the roles of two representative *silaffin* genes, *TpSil1* and *TpSil3*, in *Thalassiosira pseudonana* through gene overexpression and knockout. Due to high sequence homology, *TpSil1* and *TpSil2* were both disrupted, while the homozygous *TpSil3* knockout was proven to be lethal. Overexpression of *silaffins* increased cellular silica content, whereas the knockouts reduced silicification but enhanced cell growth and photosynthetic efficiency. TpSil3 regulates both microscale overall size and mesoscale features, while TpSil1/2 exclusively contribute to mesoscale morphology. These genetic modifications resulted in significant changes to the physicochemical and optical properties of bulk frustules, showcasing their potential in diverse applications. This study provides valuable insights into the functional roles of *silaffins* in diatoms, advancing the synthesis of tailored nanostructured silica materials through synthetic biology.

## Acknowledgements

The authors are grateful to Prof. Nils Kröger (TU Dresden) for valuable suggestions regarding the biolistic transformation of *Thalassiosira pseudonana*. The authors wish to thank Xiaofei Yan (QIBEBT) for assistance with scanning electron microscopy. The authors thank Qi An (OUC) and Keyi Jiao (OUC) for help with statistical analysis. The work was supported by the General Program of the National Natural Science Foundation of China (No. U22A20588 and No. 41976118).

