## Supplementary material for "Silaffins-Driven Genetic Engineering of Diatom Cell Walls: Insight into Biosilica Morphology and Nanomaterial Design": SI Appendix

**This file includes:**

**Supplementary Text S1 and S2**

**Figures S1 to S14**

**Tables S1 to S5**

**SI Reference** (All references are presented in the article)

**Supplementary Text**

**S1. Plasmid Constructions**

In this study, plasmids were constructed using homology-based recombination to seamlessly join the DNA fragments at their overlap regions. The general procedure is as follows: (1) Primer Design: Primers were designed with 20 base pair overlaps to adjacent DNA fragments using the NEBuilder assembly tool (nebuilder.neb.com), NCBI Primer-BLAST, and Primer Premier 5. (2) PCR Amplification: Target DNA fragments were amplified using the designed primers with FastPfu PCR Mix (TransGen, AS221). (3) Purify PCR Products: The PCR products were purified using the Gel Extraction Kit (OMEGA, D2485) to remove any impurities. (4) DNA Assembly: The purified DNA fragments were mixed and assembled using the NEBuilder® HiFi DNA Assembly Cloning Kit (E2621S, NEBuilder® HiFi Kit). (5) Transformation: The assembled recombinant plasmid was transformed into competent *E. coli* (TransGen, CD501). (6) Screening and Validation: Transformed colonies were selected, and the recombinant plasmids were verified by PCR amplification, plasmid extraction (OMEGA, D6943) and DNA sequencing (Sangon, Shanghai, CN).

It is worth noting that seamless cloning avoids the introduction of restriction enzyme sites. All experimental procedures were performed according to the manufacturers' instructions. The detailed procedures for the constructions of plasmid were as follows:

*S1.1 Construction of Plasmid for Nourseothricin Resistance*

**pFN**: *Thalassiosira pseudonana* genomic DNA (gDNA) was isolated using the

HP Plant DNA Kit (OMEGA, D2485), following the manufacturer's instructions. The promoter sequence of *fcp* LHCF9 (THAPSDRAFT_7445011) (Poulsen et al., 2006) was PCR amplified from gDNA using primers 1 and 2 (Table S1). The LHCF9 terminator was amplified from gDNA using primers 3 and 4 (Table S1). *NrsR*, which codes nourseothricin acetyltransferase conferring resistance to nourseothricin, was synthesized by Sangon Biotech (Shanghai, CN) based on the corresponding sequence from pICH47732:FCP:NAT (Addgene, No.85984) (Hopes et al., 2016). The synthetic *NrsR* sequence was used as template to amplify the homologous arm-added sequence using primers 5 and 6 (Table S1). The basic vector sequence was PCR amplified from pPha-T1(Zaslavskaia et al., 2000) using primers 7 and 8 (Table S1). The four PCR products were gel purified, then assembled into a complete plasmid using the NEBuilder® HiFi Kit, resulting in the nourseothricin resistance plasmid pTpFcpNrsR (pFN). The main steps for constructing plasmid pFN are shown in Fig. S1. With the same amplification and assembly procedures, the following plasmid schemes directly illustrate the final plasmid.

*S1.2 Construction of Plasmids for Silaffin Gene Overexpression*

Total RNA of *T. pseudonana* was extracted using the E.Z.N.A.® Total RNA Kit II (OMEGA, R6934). Subsequently, ~100 ng of RNA was reverse-transcribed into cDNA using the HiScript III 1st Strand cDNA Synthesis Kit (Vazyme, R312), following the manufacturers' instructions.

**pOESil1**: The *T. pseudonana silaffin 1* (*TpSil1*, THAPSDRAFT_11360) was PCR amplified from the cDNA using primers 9 and 10 (Table S1), incorporating a hexa-histidine tag for subsequent western blot detection. Given that the cDNA nucleotide sequence identity between *TpSil1* and *TpSil2* reaches 93.56% (Fig. S2) (Poulsen and Kröger, 2004), the PCR products were a mixture of *TpSil1* and *TpSil2* sequences. Therefore, the PCR products were gel purified and ligated into Blunt Simple Cloning Vector (TransGen, CB111), then chemically transformed into *E. coli* to isolate the *TpSil1* sequence by selecting mono-clones and DNA sequencing. Subsequently, *TpSil1* sequence was used as the template to PCR amplify the homologous arm-added *TpSil1* sequence using primers 11 and 12 (Table S1). The promoter sequence of nitrate reductase (*TpNR*) (Poulsen et al., 2006) was PCR amplified from gDNA using primers 13 and 14 (Table S1). The *TpNR* terminator region was PCR amplified from gDNA using primers 15 and 16 (Table S1). The basic vector sequence was PCR amplified from pFN using primers 17 and 18 (Table S1). After gel purification, these DNA fragments were assembled using NEBuilder® HiFi Kit, resulting in the *TpSil1* overexpression plasmid pTpFcpNrsRNRTpSil1 (pOESil1, Fig. S3A).

**pOESil3**: The *T. pseudonana silaffin 3* (*TpSil3*, THAPSDRAFT_25921) was PCR amplified from cDNA using primers 19 and 20 (Table S1) to incorporate a hexa-histidine tag. Subsequently, the PCR products were gel purified and used as the template to PCR amplify the homologous arm-added *TpSil3* sequence using primers 21 and 22 (Table S1). The basic vector sequence containing the *TpNR* expression cassette was PCR amplified from pOESil1 using primers 23 and 24 (Table S1). Both DNA fragments were gel purified and assembled by NEBuilder® HiFi Kit, resulting in the *TpSil3* overexpression plasmid pTpFcpNrsRNRTpSil3 (pOESil3, Fig. S3B).

*S1.3 sgRNA for CRISPR/Cas9-mediated Silaffins Knockout*

To generate the knockout strains of *T. pseudonana*, we employed CRISPR/Cas9 method for genome editing.

**sgRNA design**: The single guide RNAs (sgRNAs) targeting *TpSil1* and *TpSil3* were designed using CRISPOR (crispr.tefor.net) (Concordet and Haeussler, 2018) and CRISPRdirect (crispr.dbcls.jp) (Naito et al., 2015), with specificity against the *T. pseudonana* genome (strain CCMP1335, ASM14940v2). For each *silaffin* gene, three sgRNAs-encoding DNA sequences were selected (Fig. S4, Table S3).

***In vitro* sgRNA efficacy validation**: To validate the efficacy of the designed sgRNAs, we performed an *in vitro* incubation of Cas9 and sgRNA for target DNA cleavage. The sgRNAs were synthesized by Genscript (Nanjing, CN).

For targeted DNA, *TpSil1* was PCR amplified from the *T. pseudonana* gDNA using primers 25 and 26 (Table S1). The PCR products were gel purified and ligated into the Blunt Simple cloning vector, followed by chemical transformation into *E. coli* to isolate the homogenous *TpSil1* sequence through DNA sequencing. Plasmid DNA from the *TpSil1*-transformed *E. coli* strain was extracted and used as the template to PCR amplify the 5' end-extended *TpSil1* sequence using primers 26 and 29 (Table S1). Similarly, *TpSil3* was PCR amplified from the gDNA using primers 27 and 28 (Table S1). The PCR products were gel purified and ligated into Blunt Simple Cloning Vector, then chemically transformed into *E. coli*. Plasmid DNA from the DNA-sequenced strain was extracted and used as the template to PCR amplify the 5' end-extended *TpSil3* sequence using primers 28 and 29 (Table S1). Both 5’ end-extended sequences were used as the cleavage substrates.

The detailed procedures are as follows: (1) The components were added sequentially to an RNase-Free PCR tube according to Table S4. (2) The PCR tube was incubated at 37°C for 10 min in a PCR machine to allow the formation of the CRISPR/Cas9 ribonucleoprotein (RNP) complex. (3) 0.5 pmol of the cleavage substrate was added to the tube, followed by RNase-Free $H_2O$ to bring the final volume to 20 μL. (4) The PCR tube was incubated at 37°C for 30 min in a PCR machine to facilitate the cleavage reaction. (5) 1 μL of Proteinase K (20 mg $mL^{-1}$, Solarbio, P1121) was added to the tube, and the mixture was incubated at 56°C for 10 min in a PCR machine to digest any remaining Cas9 protein. (6) The cleavage product was evaluated by agarose gel electrophoresis. The molecular weight of the DNA bands was indicated using the 5K and 2K DNA Marker (TransGen, BM141 and BM101).

The agarose gel electrophoresis results from Fig. S5 suggested that both 5’ end-extended *TpSil1* and *TpSil3* were efficiently cut off only when Cas9 and sgRNA coexisted. All the designed sgRNAs could target the respective *silaffin* genes and direct Cas9 to execute cleavage with similar efficiency in each experiment. The sgRNA1 and 2 of each *silaffin* gene were selected for *in vivo* studies.

**Off-targets prediction:** To investigate whether other parts of the *T. pseudonana* genome besides the *silaffin* gene can be affected in the knockout mutants, potential off-target sites for *silaffin s*gRNAs were analyzed using the CRISPOR (crispr.tefor.net)

(Concordet and Haeussler, 2018). CRISPOR predicted that no binding sites with less than four mismatches for either *TpSil1* sgRNA1 or sgRNA2 are present in the *T. pseudonana* genome (Table S5). But in fact, *TpSil1*_sgRNA1 also targets *TpSil2* due to the high sequence homology between *TpSil1* and *TpSil2*. Off-target analysis did not identify this because only one corresponding gene for *TpSil1* and *TpSil2*, THAPSDRAFT_11360, is listed in the *T. pseudonana* genome database. With four mismatches allowed, there are no additional binding sites for *TpSil3* sgRNA1 and sgRNA2 other than to the *TpSil3* gene (Table S5). In this study, DNA fragments of the potential off-target sites for mismatched sgRNAs were not analyzed, because previous studies demonstrated that potential off-target sites with four or more mismatches were not affected by Cas9 in both diatoms *Phaeodactylum tricornutum* (Stukenberg et al., 2018) and *T. pseudonana* (Görlich et al., 2019).

*S1.4 Construction of Plasmids for Silaffin Gene Knockout*

Domesticated human codon bias Cas9 from *Streptococcus pyogenes* with SV40 nuclear localisation signal (NLS) and enhanced yellow fluorescent protein (EYFP) were PCR-amplified from the plasmid pICH47742:FCP:Cas9YFP (Addgene No.85986) (Hopes et al., 2016) using primers 30 and 31 (Table S1) whose product was named Cas9-YFP. DNA sequence of Ori, *AmpR*, and *TpNR* promoter was PCR-amplified from plasmid pOESil (both pOESil1 and pOESil3 are feasible) using primers 32 and 33 (Table S1) whose product was named pOESil part 1. *TpNR* terminator, *TpFcp* promoter, *NrsR*, and *TpFcp* terminator were PCR-amplified from plasmid pOESil using primers 34 and 35 (Table S1) whose product was named pOESil part 2. The two sgRNAs are each placed under the *T. pseudonana* U6 promoter (Hopes et al., 2016) and the sgRNA scaffold, of which the descriptions are shown in Fig. S6. Sequences of TpSil1 U6_sgRNA1_sgRNA2 and TpSil3 U6_sgRNA1_sgRNA2 were synthesized by Genscript (Nanjing, CN).

**pKOSil1**: TpSil1 U6_sgRNA1_sgRNA2 was used as template to PCR-amplify the homologous arm-added sequences using primers 36 and 37 (Table S1). After gel purification, Cas9-YFP, pOESil part 1, pOESil part 2, and arm-added TpSil1 U6_sgRNA1_sgRNA2 were assembled using NEBuilder® HiFi Kit, resulting in the *TpSil1* gene knockout plasmid pTpFcpNrsRNRCas9YFPU6Sil1_sgRNA1_sgRNA2 (pKOSil1, Fig. S7A).

**pKOSil3**: TpSil3 U6_sgRNA1_sgRNA2 was used as template to PCR-amplify the homologous arm-added sequences using primers 36 and 37 (Table S1). Similarly, after gel purification, Cas9-YFP, pOESil part 1, pOESil part 2, and armed-added TpSil3 U6_sgRNA1_sgRNA2 were assembled, resulting in the *TpSil3* gene knockout plasmid pTpFcpNrsRNRCas9YFPU6Sil3_sgRNA1_sgRNA2 (pKOSil3, Fig. S7B).

### S2. Genomic Screening of Genetically Transformed Strains of *T. pseudonana*

Above plasmids were introduced into *T. pseudonana* using biolistic bombardment, followed by plating transformed cells on agar plates containing nourseothricin. For positive transformed strains screen, resistant clones grown from the selective agar plates were confirmed by PCR amplification, gel electrophoresis and DNA sequencing. Plant Direct PCR Kit (Vazyme, PD105) was used to isolate transformant's gDNA, following the manufacturer's instructions.

*S2.1 Screening of TpSil1 and TpSil3 overexpression strains*

*TpSil1* overexpression strains were screened using primers 1 and 2 (Table S2) to PCR amplify the specific sequence of pOESil1 (Fig. S8A). while *TpSil3* strains were screened with primers 3 and 4 (Table S2) to PCR amplify the specific sequence of pOESil3 (Fig. S8B). The expected band size was observed in genetically transformed strains, while no amplification was observed in wild-type strain, confirming the genomic integration of overexpression plasmid in transformed strains.

*S2.2 Screening of TpSil1 and TpSil3 Knockout Strains*

For selection of knockout strains, band-shift PCR was performed. Knockout strains were identified by the presence of a shorter band on the agarose gel.

*S2.2.1 Both TpSil1 and TpSil2 Knockout Strains*

For selection of *TpSil1* knockout strains, band-shift PCR was performed using primers 5 and 6 (Table S2), and several heterozygous *TpSil1* knockout strains were identified during preliminary screening (Fig. S9A). Since *T. pseudonana* is diploid, primary colonies often exhibited mosaicism, containing both wild-type and mutant alleles. After cultivation of the heterozygous strains to the exponential phase, ~$10^3$ cells were collected and re-plated onto selective plates. The sub-clones were further analyzed by PCR amplification and DNA sequencing to select homozygous knockout strains.

Due to the high sequence identity between *TpSil1* and *TpSil2* (Fig. S2), PCR amplification using primers 5 and 6 resulted in products from both *TpSil1* and *TpSil2*, complicating the identification of homozygous knockouts. To overcome this, annealing temperature-specific primers, KOSil1 Screen (Band Deletion) (Table S2, Fig. S9B) were designed. At the annealing temperature of 46°C, both *TpSil1* and *TpSil2* were PCR amplified, whereas at 58°C, only *TpSil1* was amplified (Fig. S9C). The annealing temperature for the KOSil1 Screen (Band Deletion) primers was optimized. Sub-clones that produced bands at 46°C but showed no amplification at 58°C were identified as homozygous *TpSil1* knockout strains. Among the 30 sub-clones analyzed, 7 exhibited a homozygous *TpSil1* knockout pattern (Fig. S9D). These sub-clones were then PCR amplified using primers 5 and 6 (Table S2) and transformed into *E. coli* for DNA sequencing. At least 5 colonies per *E. coli* transformant were sequenced, and a sub-clone was confirmed as homozygous knockout only if sequencing results exclusively verified the knockout of the *TpSil1* sequence. However, sequencing results not only confirmed the knockout of *TpSil1* sequence, but also the *TpSil2* sequence. Due to the high sequence similarity between the two genes, *TpSil1*_sgRNA1 also targeted *TpSil2*, resulting in the disruption of both genes. Since the Cas9 and *TpSil1*_sgRNA1 will continue to be expressed in the transformed strains, leading to the persistent targeting of *TpSil2*, two sub-clones were selected, designated KO1/2-1 and KO1/2-2, in which both *TpSil1* and *TpSil2* were knocked out for further investigation. DNA sequencing results for KO1/2-1 and KO1/2-2 (Fig. S11A) confirmed that the disrupted *TpSil1* and *TpSil2* genes encode significantly truncated polypeptides (Fig. S12).

*S2.2.2 Incomplete TpSil3 knockout strains*

Band-shift PCR using primers 9 and 10 (Table S2) was employed to screen for homozygous *TpSil3* knockout strains. During the process, two heterozygous knockout strains were identified in the initial analysis (Fig. S10A) and were re-plated. The number of sub-clones obtained was significantly lower than for *TpSil1.* A total of 168 sub-clones were collected and analyzed by PCR amplification with primers 9 and 10, among which only three strains exhibited a homozygous knockout pattern on the gel (Fig. S10B). DNA sequencing confirmed that two of these strains shared the same knockout site, while another strain exhibited a different mutation pattern. Notably, in these strains, there was always one *TpSil3* allele showed a base deletion in multiples of three (Fig. S11B), resulting in the deletion of a small number of amino acids (Fig. S12). Sequencing of 20 additional sub-clones without a homozygous knockout pattern on the gel revealed the presence of a complete *TpSil3* allele in all cases. Consequently, only two incomplete *TpSil3* knockout strains were identified and designated as iKO3-1 and iKO3-2.

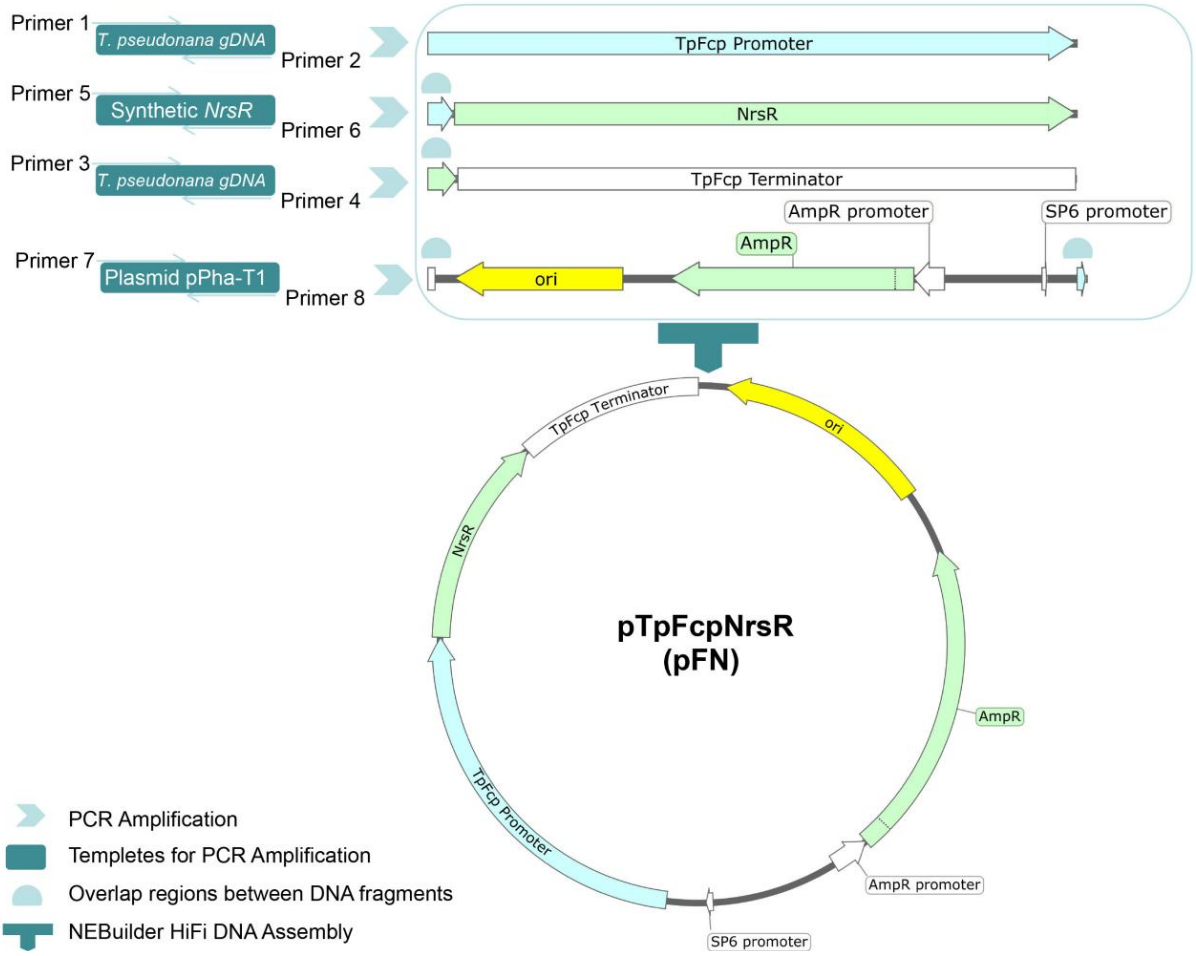


Fig. S1. Scheme of the pTpFcpNrsR (pFN) plasmid construction designed for nourseothricin resistance in *T. pseudonana*. TpFcp: *T. pseudonana* fucoxanthin chlorophyll a/c binding protein LHCF9. NrsR: nourseothricin acetyltransferase which confers resistance to nourseothricin. ori: origin of replication. AmpR: β-lactamase which confers resistance to ampicillin. Primers are listed in Table S1.

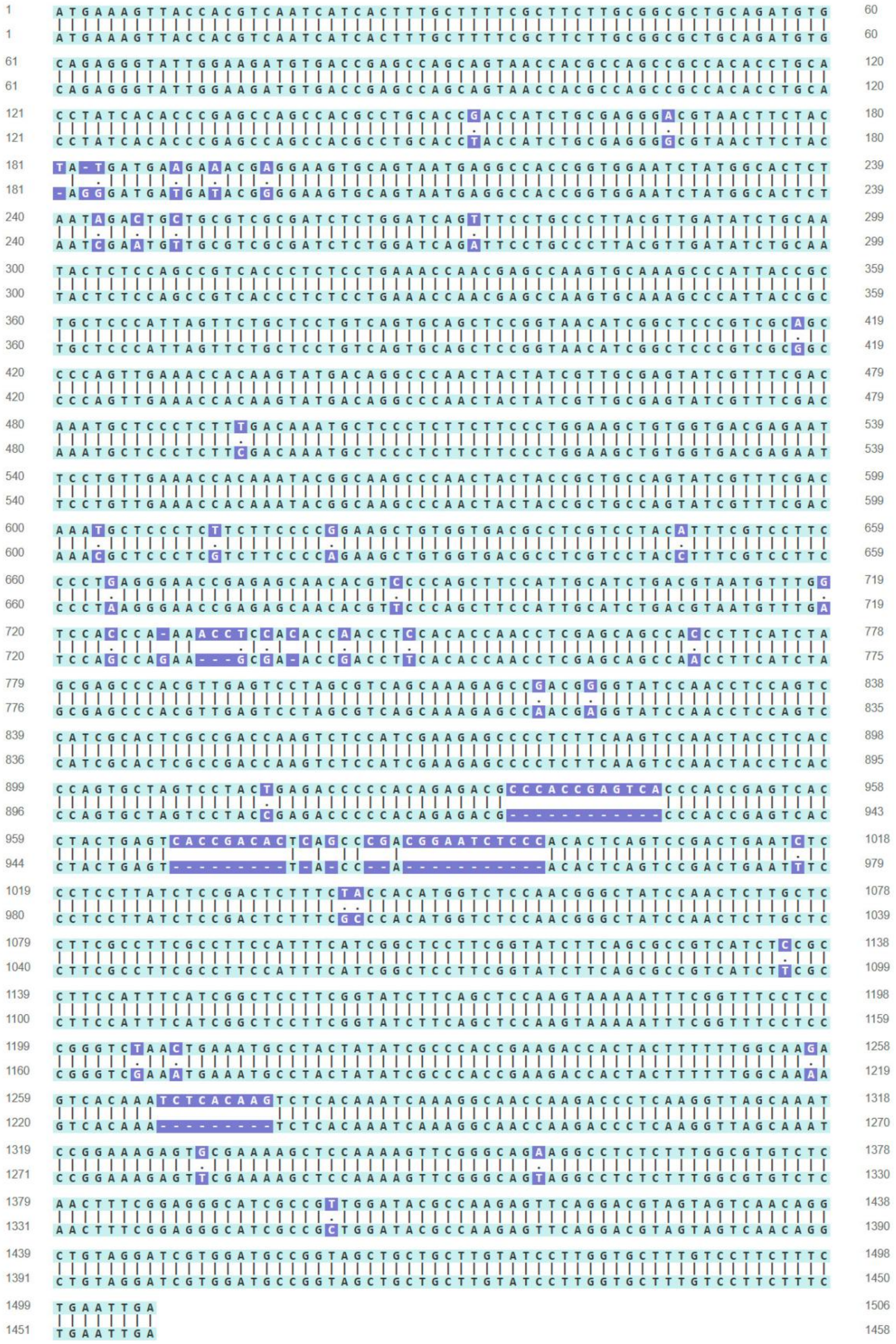

Fig. S2. cDNA nucleotide sequence identity analysis between *TpSil1* (upper) and *TpSil2* (lower).

(A)

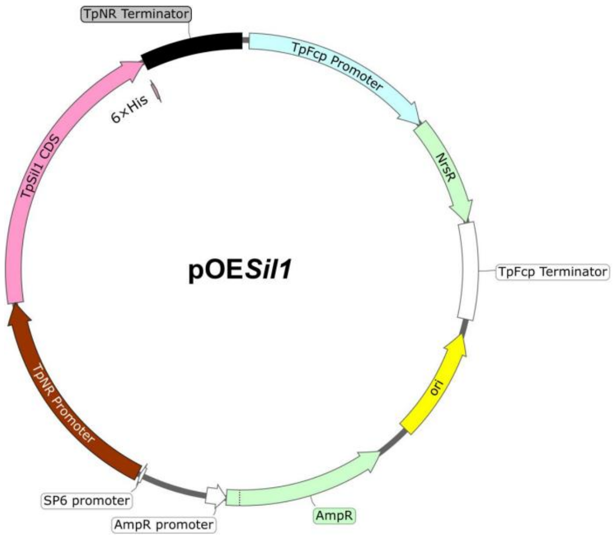


(B)

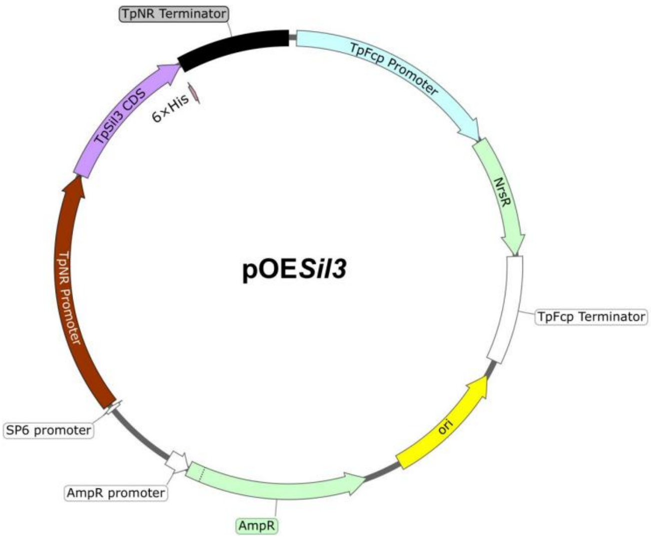


Fig. S3. Scheme of the plasmids designed for overexpression of *silaffin* genes in *T. pseudonana.* TpFcp: *T. pseudonana* fucoxanthin chlorophyll a/c binding protein LHCF9. NrsR: nourseothricin acetyltransferase which confers resistance to nourseothricin. ori: origin of replication, AmpR: β-lactamase which confers resistance to ampicillin. TpNR: *T. pseudonana* nitrate reductase. (A) *TpSil1* overexpression plasmid pOESil1 and (B) *TpSil3* overexpression plasmid pOESil3.

(A)

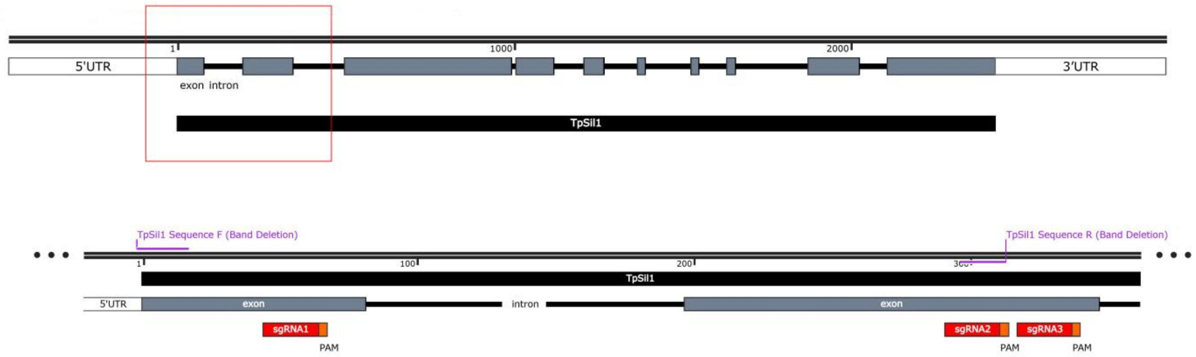


(B)

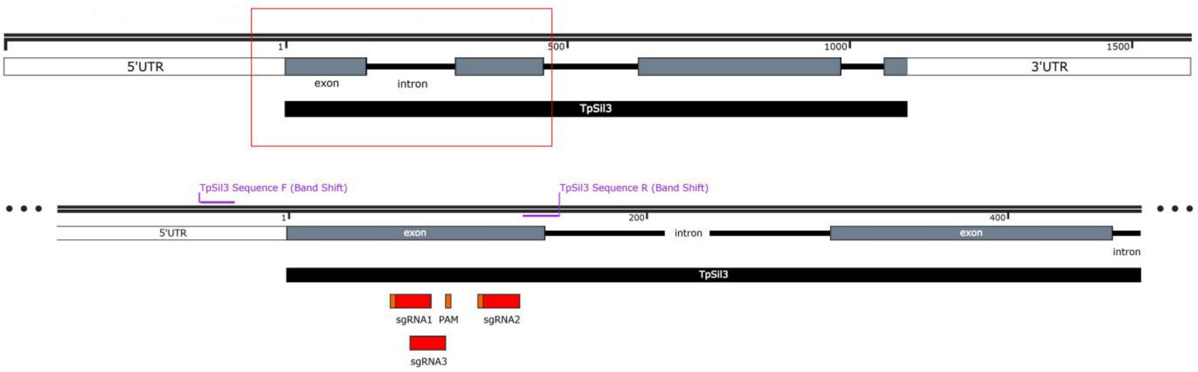


Fig. S4. Schematic representation of the binding sites of sgRNAs to *TpSil1* (A) and *TpSil3* (B).

(A)

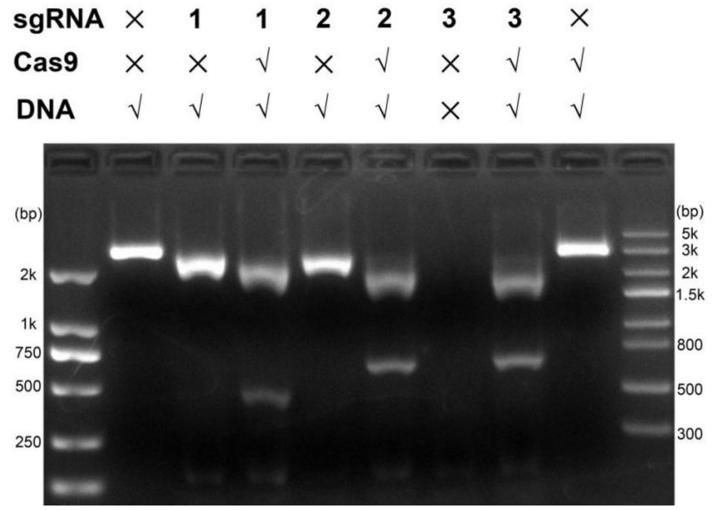


(B)

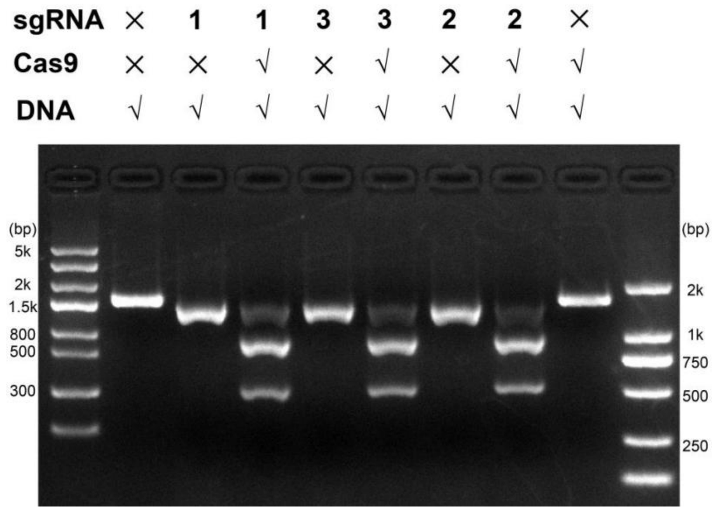


Fig. S5. *In vitro* cleavage of Cas9 and sgRNA for 5' end-extended *TpSil1* (A) and *TpSil3* (B). √ or × indicates the presence or absence of the substance.

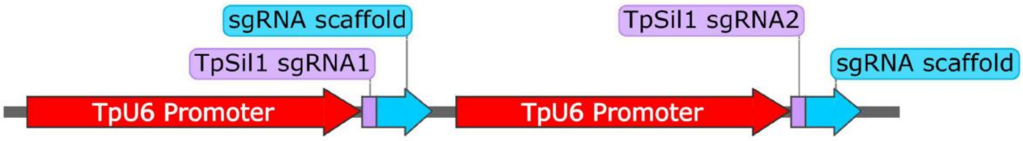


GCTACTAGAATTCGAGCTCGGAGCTTCATCAAGAGAGCAACCAACAAAGAAGAGGCGAACACGGTGCACAGCACCGTCTGCATACTTCCG
ATTTGCAATTCACAACATATGATACATCATCCATCTCTCTGAGAAAACTGGGCACTGTTAGAATCTGGGATTTATCAACAGCCACCAACTAT
CTACTCAACGCAATTGGGGCACTTGATAATGTGACAATCCTCCTTGCTCCTTTCCTTGTCGCCTCAAACAACGTCGGGAACTGCACAGAAA
CATCGCTACTTATTCACGCCTCACCTCTCTTTCCTCCTTTACTCACTTATTCGCGCCTCACCTCTCTTTCCTCCTTTACTCAAGTCCTCTGCT
TGCCGGGTTTATTCTTCCGTTTGTGTTGTTGCGGTCTGATAGGATTTGTTTGGAGACGAACCTTTTTTTTTGATTGCTTTATAAATTAACGTT
TTGCCGAAATTGTCGGCGCTGCAGATGTGCAGAGTTTTAGAGCTAGAAATAGCAAGTTAAAATAAGGCTAGTCCGTTATCAACTTGAAAAA
GTGGCACCGAGTCGGTGCTTTTTTCGCTTTACGAATTCCCATGGGGAGCTTCATCAAGAGAGCAACCAACAAAGAAGAGGCGAACACGGT
GCACAGCACCGTCTGCATACTTCCGATTTGCAATTCACAACATATGATACATCATCCATCTCTCTGAGAAAACTGGGCACTGTTAGAATCTG
GGATTTATCAACAGCCACCAACTATCTACTCAACGCAATTGGGGCACTTGATAATGTGACAATCCTCCTTGCTCCTTTCCTTGTCGCCTCAA
ACAACGTCGGGAACTGCACAGAAACATCGCTACTTATTCACGCCTCACCTCTCTTTCCTCCTTTACTCACTTATTCGCGCCTCACCTCTCTT
TCCTCCTTTACTCAAGTCCTCTGCTTGCCGGGTTTATTCTTCCGTTTGTGTTGTTGCGGTCTGATAGGATTTGTTTGGAGACGAACCTTTTT
TTTTGATTGCTTTATAAATTAACGTTTTGCCGAAATTGTTCTACTATGATGAAGAAACGGTTTTAGAGCTAGAAATAGCAAGTTAAAATAAGG
CTAGTCCGTTATCAACTTGAAAAAGTGGCACCGAGTCGGTGCTTTTTTCGCTCAGAGAGGATGCACATGTGACCGA

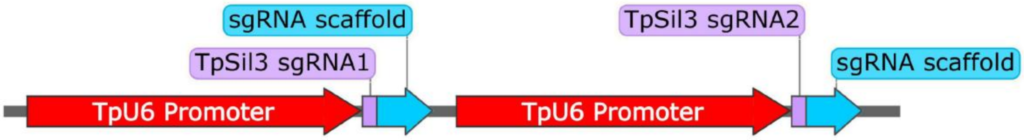


GCTACTAGAATTCGAGCTCGGAGCTTCATCAAGAGAGCAACCAACAAAGAAGAGGCGAACACGGTGCACAGCACCGTCTGCATACTTCCG
ATTTGCAATTCACAACATATGATACATCATCCATCTCTCTGAGAAAACTGGGCACTGTTAGAATCTGGGATTTATCAACAGCCACCAACTAT
CTACTCAACGCAATTGGGGCACTTGATAATGTGACAATCCTCCTTGCTCCTTTCCTTGTCGCCTCAAACAACGTCGGGAACTGCACAGAAA
CATCGCTACTTATTCACGCCTCACCTCTCTTTCCTCCTTTACTCACTTATTCGCGCCTCACCTCTCTTTCCTCCTTTACTCAAGTCCTCTGCT
TGCCGGGTTTATTCTTCCGTTTGTGTTGTTGCGGTCTGATAGGATTTGTTTGGAGACGAACCTTTTTTTTTGATTGCTTTATAAATTAACGTT
TTGCCGAAATTGTTCAAGAGTTCTCAATCGGCGGTTTTAGAGCTAGAAATAGCAAGTTAAAATAAGGCTAGTCCGTTATCAACTTGAAAAAG
TGGCACCGAGTCGGTGCTTTTTTCGCTTTACGAATTCCCATGGGGAGCTTCATCAAGAGAGCAACCAACAAAGAAGAGGCGAACACGGTG
CACAGCACCGTCTGCATACTTCCGATTTGCAATTCACAACATATGATACATCATCCATCTCTCTGAGAAAACTGGGCACTGTTAGAATCTGG
GATTTATCAACAGCCACCAACTATCTACTCAACGCAATTGGGGCACTTGATAATGTGACAATCCTCCTTGCTCCTTTCCTTGTCGCCTCAAA
CAACGTCGGGAACTGCACAGAAACATCGCTACTTATTCACGCCTCACCTCTCTTTCCTCCTTTACTCACTTATTCGCGCCTCACCTCTCTTT
CCTCCTTTACTCAAGTCCTCTGCTTGCCGGGTTTATTCTTCCGTTTGTGTTGTTGCGGTCTGATAGGATTTGTTTGGAGACGAACCTTTTTT
TTTGATTGCTTTATAAATTAACGTTTTGCCGAAATTGTTTTCGAGCTGTGCATGGACAGTTTTAGAGCTAGAAATAGCAAGTTAAAATAAGGC
TAGTCCGTTATCAACTTGAAAAAGTGGCACCGAGTCGGTGCTTTTTTCGCTCAGAGAGGATGCACATGTGACCGA

Fig. S6. Descriptions of the TpSil1 U6_sgRNA1_sgRNA2 (upper) and TpSil3 U6_sgRNA1_sgRNA2 (lower) constructs. *T. pseudonana* U6 promoter sequences are shown in red; the sgRNA sequences are shown in pinkish purple; the sgRNA scaffold sequences are shown in turquoise.

(A)

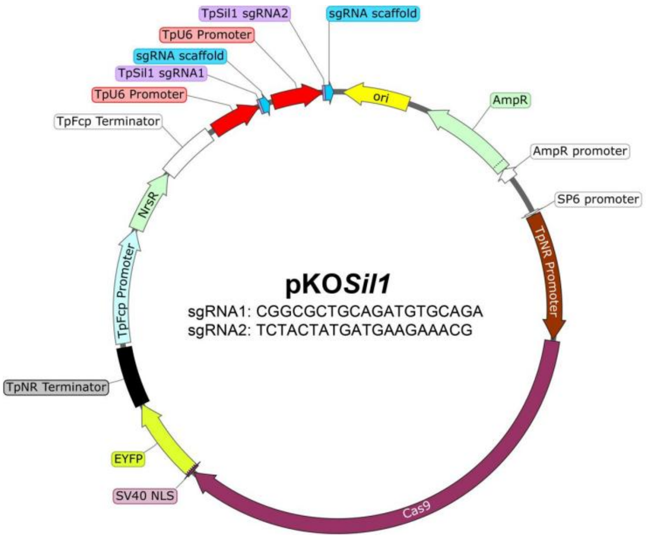


(B)

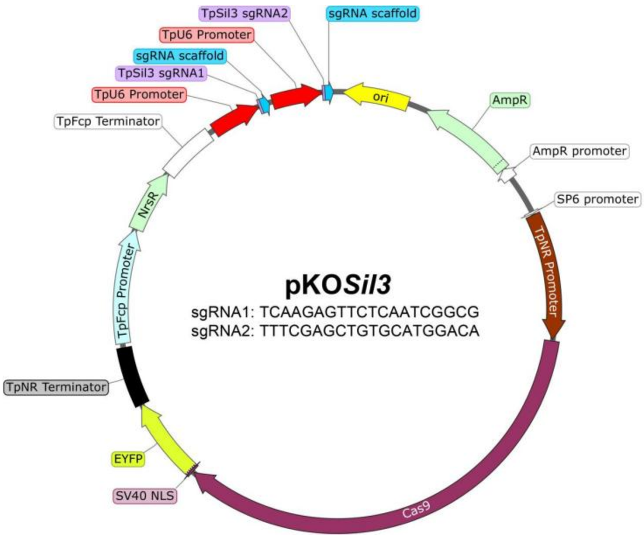


Fig. S7. Scheme of the plasmids designed for CRISPR/Cas9-mediated knockout of *silaffin* genes in *T. pseudonana*. TpFcp: *T. pseudonana* fucoxanthin chlorophyll a/c binding protein LHCF9. NrsR: nourseothricin acetyltransferase which confers resistance to nourseothricin. ori: origin of replication. AmpR: β-lactamase which confers resistance to ampicillin. TpNR: *T. pseudonana* nitrate reductase. Cas9: Cas9 nuclease. SV40 NLS: simian virus 40 nuclear localization signal. EYFP: enhanced yellow fluorescent protein. In the center of the scheme, the sgRNA sequences specific the indicated *silaffin* gene are shown. (A) *TpSil1* knockout plasmid pKOSil1 and (B) *TpSil3* knockout plasmid pKOSil3.

(A)

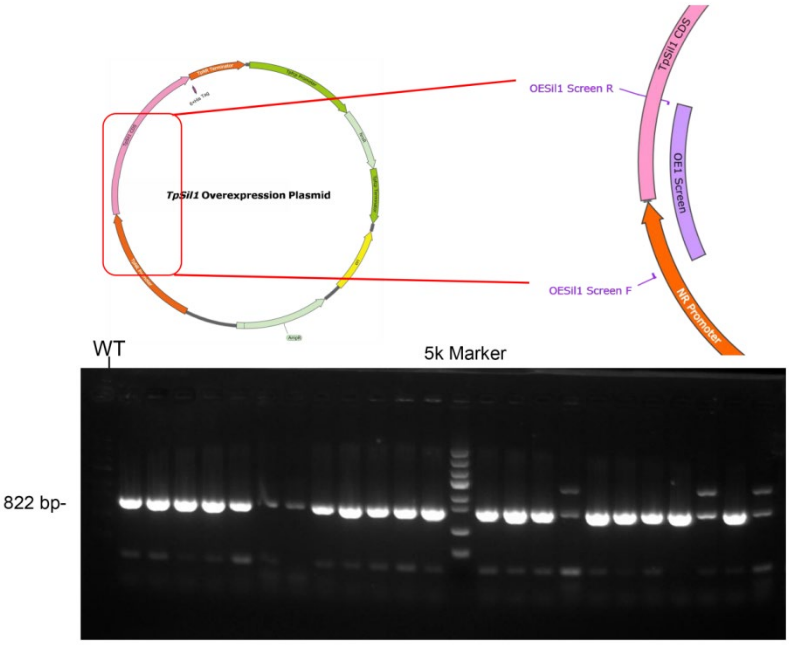


(B)

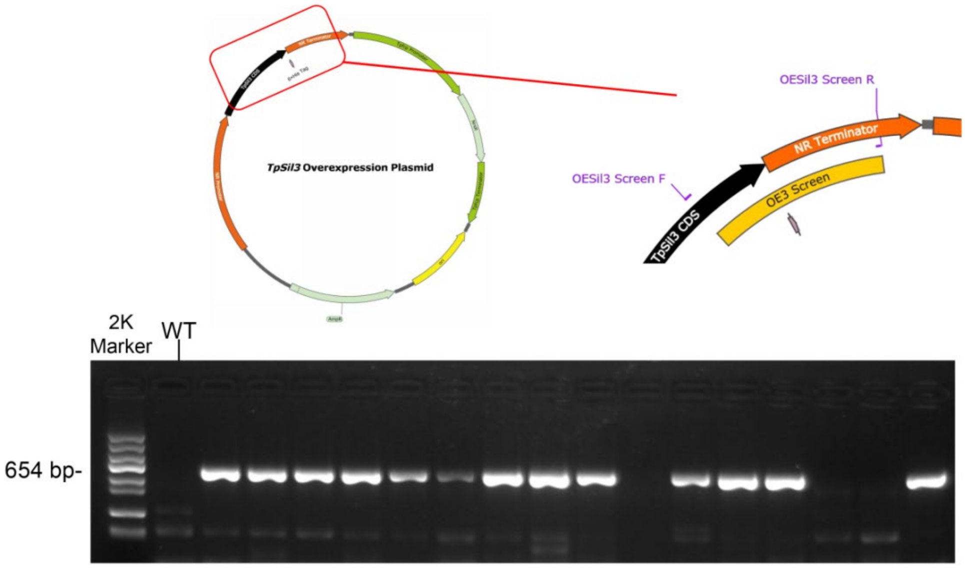


Fig. S8. Genomic integration verification of overexpression plasmid in transformed strains by PCR amplification and gel electrophoresis. (A) Screening of *TpSil1* overexpression strains using primers 1 and 2 (Table S2) that specific to pOESil1; (B) Screening of *TpSil3* overexpression strains using primers 3 and 4 (Table S2) that specific to pOESil3.

(A)

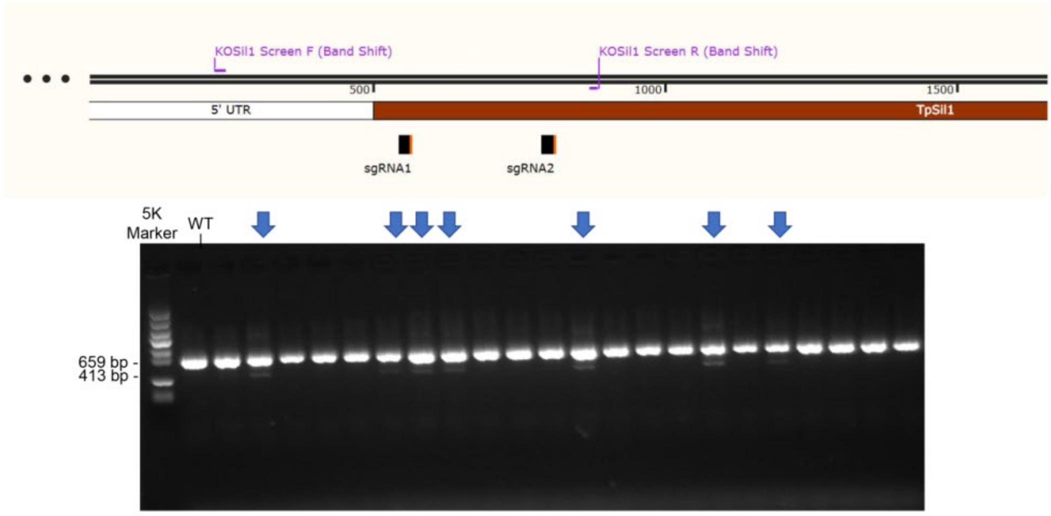


(B)

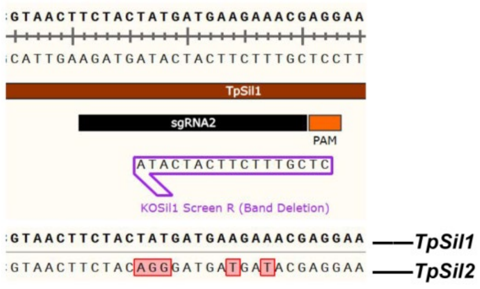


Fig. S9. Screening of homozygous *TpSil1* knockout strains. (A) Preliminary screening of heterozygous *TpSil1* knockout strains by band-shift PCR amplification using primers 5 and 6 (Table S2). Potential knockout strains are highlighted by blue arrows. (B) The KOSil1 Screen (Band Deletion) primers (Table S2), with the reverse primer containing 5 mismatches to *TpSil2*.

(C)

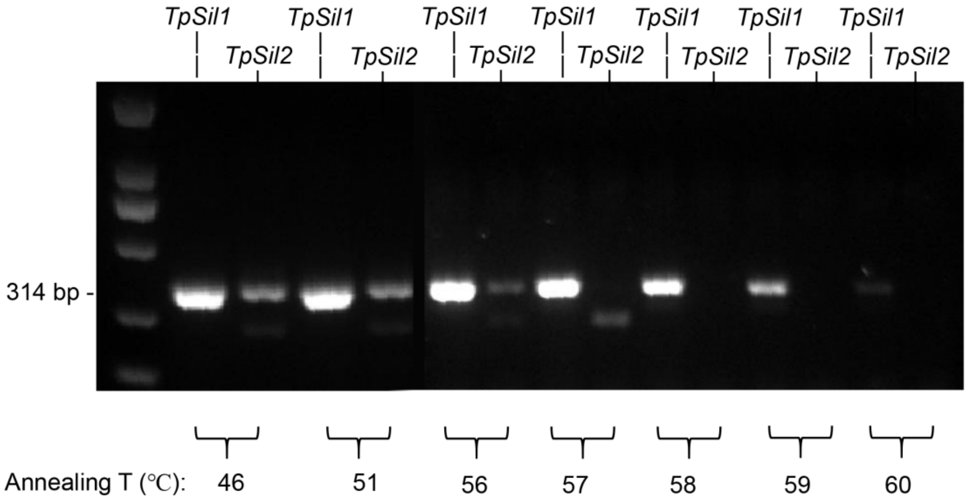


(D)

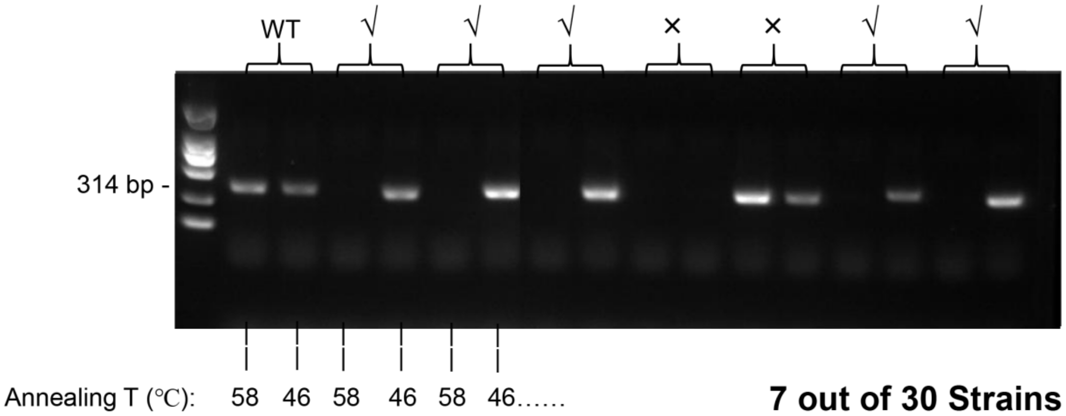


Fig. S9 (continued). (C) Optimization of PCR annealing temperature for KOSil1 Screen (Band Deletion) primers. (D) Screening of homozygous *TpSil1* knockout strains from the re-plated sub-clones using KOSil1 Screen (Band Deletion) primers. Sub-clones that produced bands at 46℃ but showed no amplification at 58℃ were identified as homozygous *TpSil1* knockout strains.

(A)

(B)

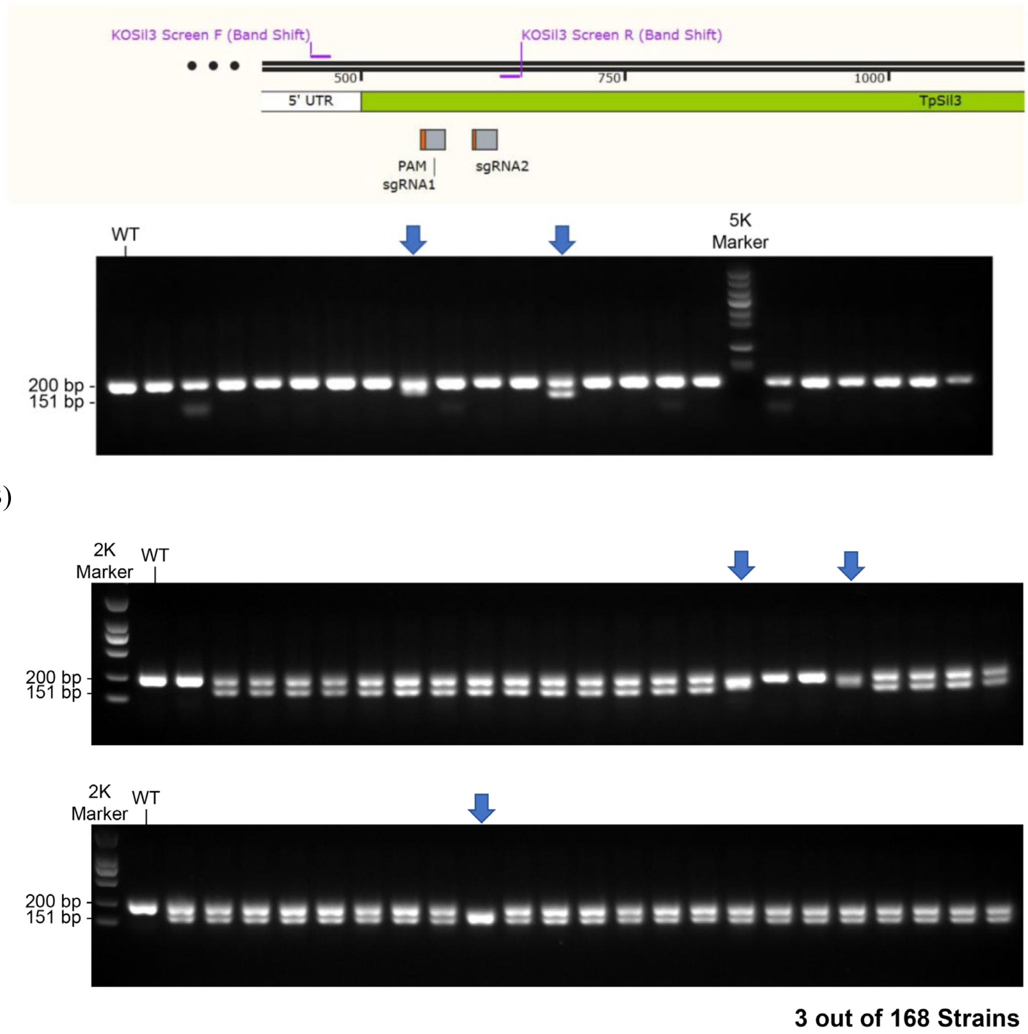


Fig. S10. Screening of homozygous *TpSil3* knockout strains. (A) Preliminary screening of *TpSil3* knockout strains by band-shift PCR amplification using primers 9 and 10 (Table S2). (B) Screening of homozygous *TpSil3* knockout strains from the re-plated sub-clones using the same primers. Potential knockout strains are highlighted by blue arrows.

(A)

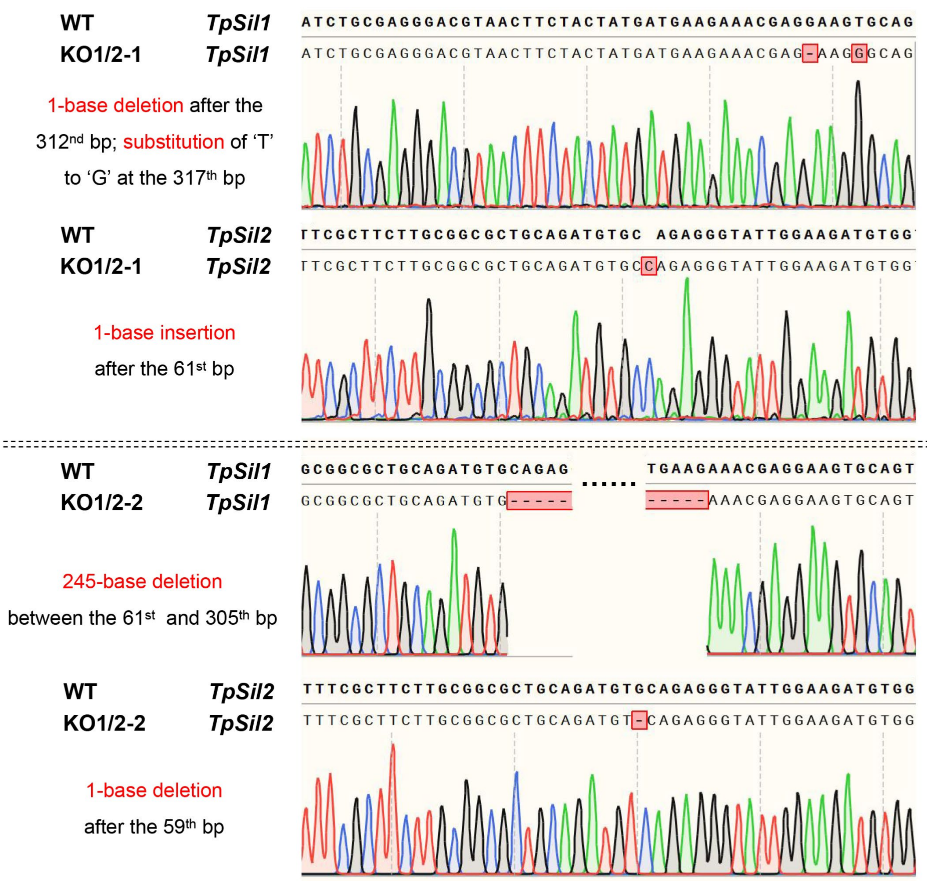


Fig. S11. DNA sequencing results of the *silaffins* knockout strains. The DNA sequences of the edited regions, the editing strategy, sequencing chromatogram, and the corresponding wild-type sequences are shown for each knockout strains. (A) Knockout strains KO1/2-1 and KO1/2-2 with edited *TpSil1* and *TpSil2* genes.

(B)

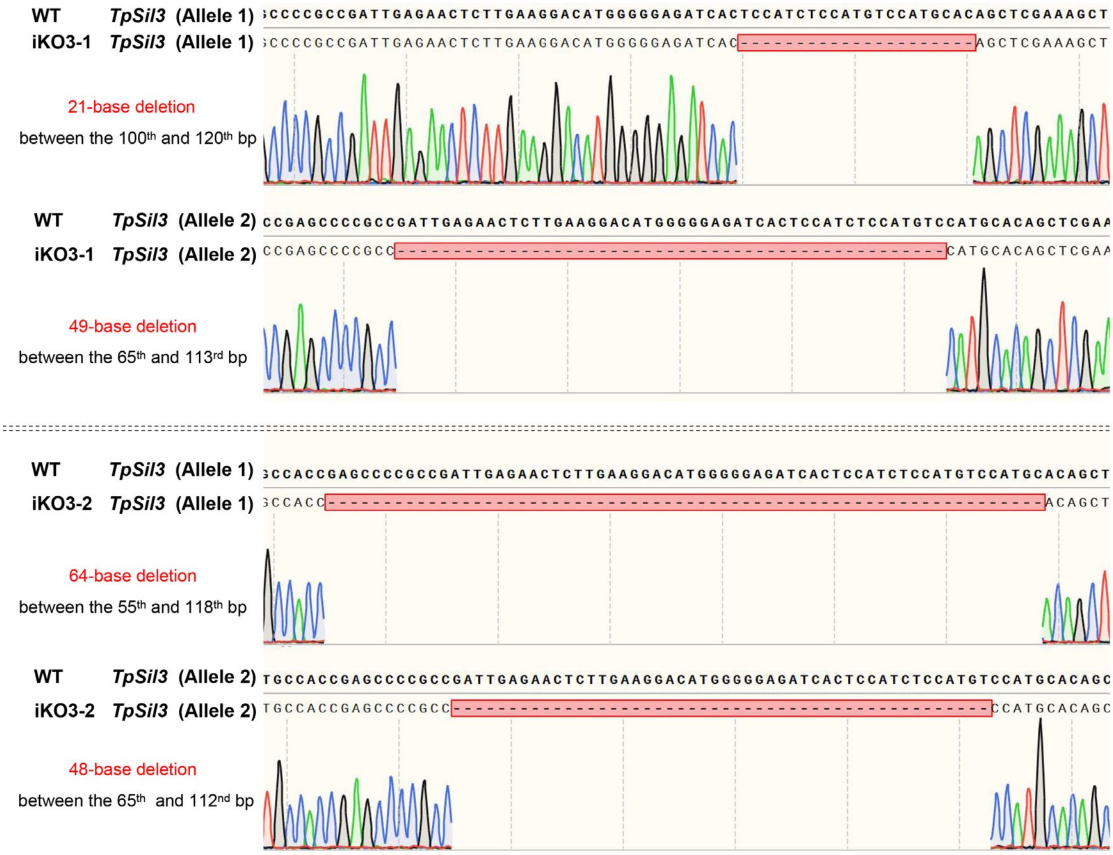


Fig. S11 (continued). (B) Incomplete knockout strains iKO3-1 and iKO3-2 with two edited *TpSil3* alleles.

**KO1/2-1** ***TpSil1***
MKVTTSIITLLFASCGAADVQRVLEDVTEPAVTTPAATPAPITPEPATPAPTICEGRNFYYDEETR RAVMRPPVESMAL*
**KO1/2-1** ***TpSil2***
MKVTTSIITLLFASCGAADV PEGIGRCDRASSNHASRHTCTYHTRASHACTYHLRGA*

**KO1/2-2** ***TpSil1***
MKVTTSIITLLFASCGAAD EKRGSAVMRPPVESMAL*
**KO1/2-2** ***TpSil2***
MKVTTSIITLLFASCGAADV RGYWKM*

**iKO3-1** ***TpSil3*** **(Allele 1)**
MKTSAIALLAVLATTAATEPRRLRTLEGHGGDHSSKAEKQAIEAAVEEDVAGPAKAAKLFKPKASKAGSMPDEAGAKSAKMSMDTKSG
KSEDAAAVDAKASKESHMSISGDMSMAKSHKAEAEDVTEMSMAKAGKDEASTEDMCMPFAKSDKEMSVKSKQGKTEMSVADAKAS
KESSMPSSKAAKIFKGKSGKSGSLSMLKSEKASSAHSLSMPKAEKVHSMSA*
(Deletion of 'ISMSMHS' between wild type p35 and p41)
**iKO3-1** ***TpSil3*** **(Allele 2)**
MKTSAIALLAVLATTAATEPR PCTARKLRSKPSRQLLRRMLLALQRQPSFSSPKQARLVPCLMRPVQRVPR*

**iKO3-2** ***TpSil3*** **(Allele 1)**
MKTSAIALLAVLATTAAT TARKLRSKPSRQLLRRMLLALQRQPSFSSPKQARLVPCLMRPVQRVPR*
**iKO3-2** ***TpSil3*** **(Allele 2)**
MKTSAIALLAVLATTAATEPRPMHSSKAEKQAIEAAVEEDVAGPAKAAKLFKPKASKAGSMPDEAGAKSAKMSMDTKSGKSEDAAAVD
AKASKESHMSISGDMSMAKSHKAEAEDVTEMSMAKAGKDEASTEDMCMPFAKSDKEMSVKSKQGKTEMSVADAKASKESSMPSSK
AAKIFKGKSGKSGSLSMLKSEKASSAHSLSMPKAEKVHSMSA*
(Substution of 'LRTLEGHGGDHSISMS' to 'P' between wild type p22 and p38)

Fig. S12 Amino acid sequence encoded by the mutated *silaffin* genes in the knockout strains. Differences to the wild-type sequence are highlighted in red. Termination of the amino acid sequence is indicated by an asterisk.

(A)

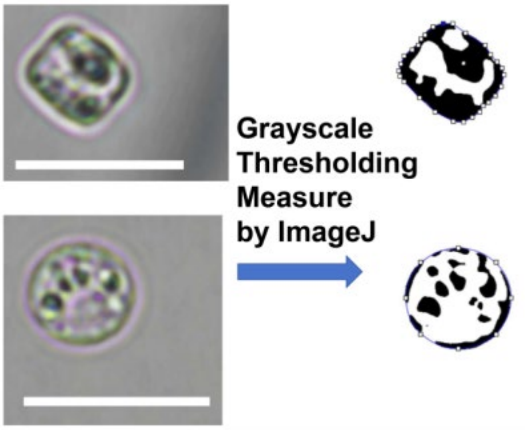


(B)

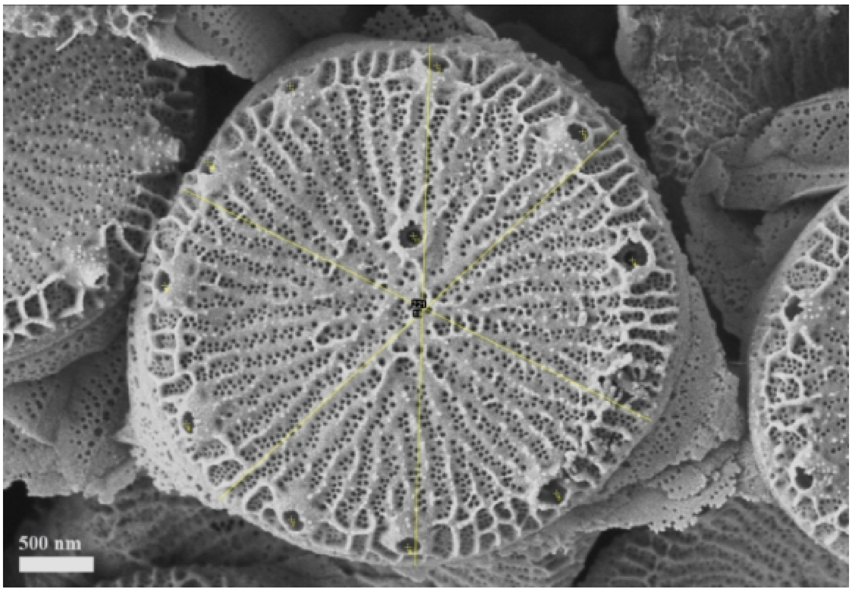


Fig. S13. (A) Identification of cell dimensions from light microscopy images. Upper panel: cylindrical cell profile; lower panel: circular cell profile. Scale bar: 10 µm. (B) Measurement of valve diameter and fultoportula number of frustule valve from a scanning electron microscopy image.

(A)

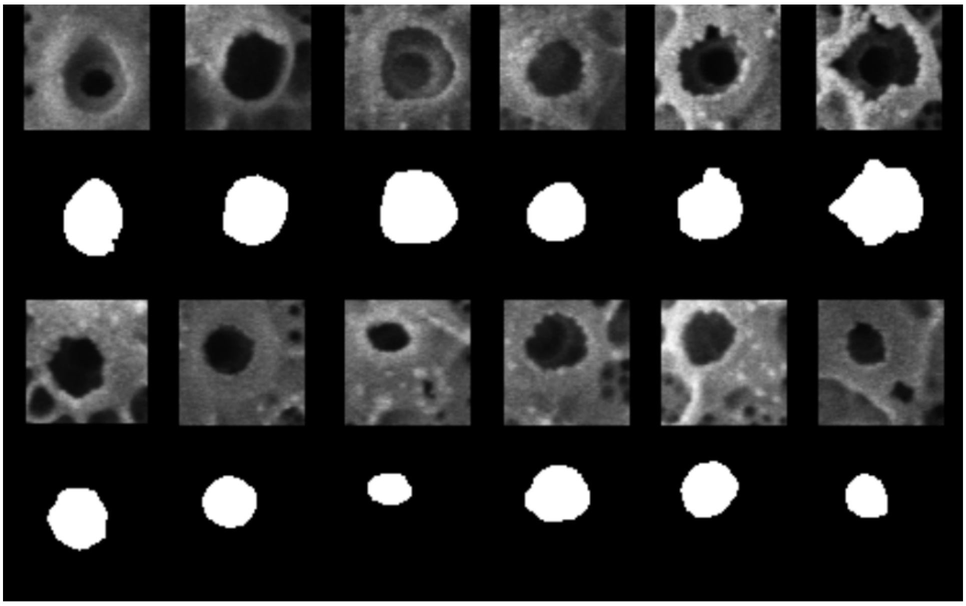

(B)

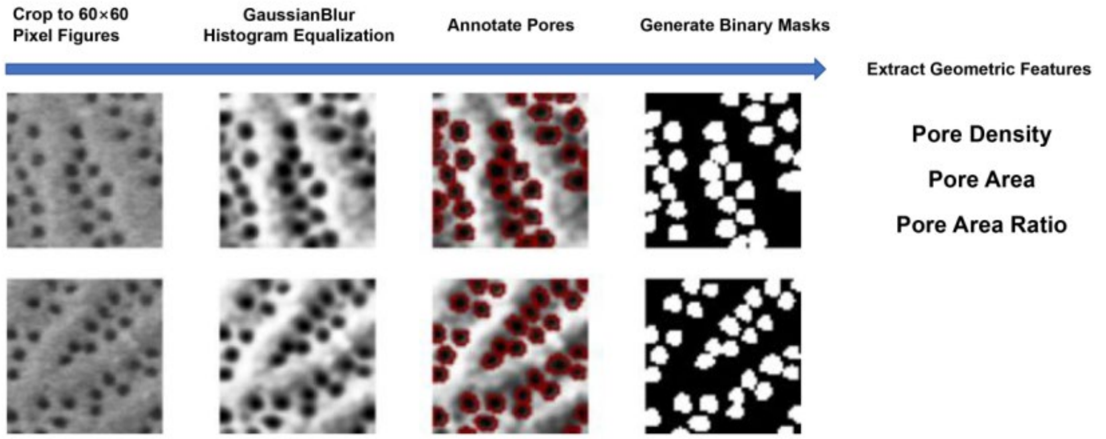


**Fig. S14.** Analysis of fultoportula area (A) and cribrum pore patterns (B) of the wild-type frustule within a 60 × 60-pixel field from scanning electron microscopy images.

**Table S1.** Oligonucleotides for plasmid constructions.

| Name | Sequence | Primer No. |
|---|---|---|
| pFN *TpFCP* Promoter F | CAAGGACGGCATCCATGGCA | 1 |
| pFN *TpFCP* Promoter R | TTTGGTATTGGTTTGGTAAATCAGATAATTTATGAGAGG | 2 |
| pFN *TpFCP* Terminator F | *CTACATGAGCATGCCCTGCCCCTGA*ATACTGGATTGGTGAATCAATGAGC | 3 |
| pFN *TpFCP* Terminator R | GGGAGAACTGGAGCAGCTAC | 4 |
| pFN *NrsR* F | *TCTGATTTACCAAACCAATACCAAA*ATGACCACTCTTGACGACACGGC | 5 |
| pFN *NrsR* R | TCAGGGGCAGGGCATGCTCA | 6 |
| pFN pPha-T1 Part F | *AAGTAGTAGCTGCTCCAGTTCTCCC*AACGCAGGAAAGAACATGTG | 7 |
| pFN pPha-T1 Part R | *CGTTTTGCCATGGATGCCGTCCTTG*TAATCTGAATCTGATCCTCCTTTTC | 8 |
| *TpSil1* (6×his) F | ATGAAAGTTACCACGTCAATCATCACTTTGC | 9 |
| *TpSil1* (6×his) R | TCA**ATGATGATGATGATGATG**ATTCAGAAAGAAGGAC | 10 |
| pOESi1 *TpSil1* F | *TATCATAATC*ATGAAAGTTACCACGTCAATC | 11 |
| pOESi1 *TpSil1* R | *AGCATCCTCA*TCAATGATGATGATGATGATG | 12 |
| pOESi1 *TpNR* Promoter F | *CAGATCCCCC*CCCTACATCATTGACGGATC | 13 |
| pOESi1 *TpNR* Promoter R | *TAACTTTCAT*GATTATGATAGTTGTTGGTTCC | 14 |
| pOESi1 *TpNR* Terminator F | *TCATCATTGA*TGAGGATGCTCATCGTTTTG | 15 |
| pOESi1 *TpNR* Terminator R | *GGTCGCTCGA*GGATGCATTGCCTGATAATATC | 16 |
| pOESi1 pFN part F | *CAATGCATCC*TCGAGCGACCATGGAAAAGG | 17 |
| pOESi1 pFN part R | *TGATGTAGGG*GGGGGATCTGGTTCTATAGTG | 18 |
| *TpSil3* (6×his) F | ATGAAGACTTCTGCCATTGCATTGCTTGC | 19 |
| *TpSil3* (6×his) R | TCA**ATGATGATGATGATGATG**AGCGCTCATGGAGTG | 20 |
| pOESil3 *TpSil3* F | *TATCATAATC*ATGAAGACTTCTGCCATTG | 21 |
| pOESil3 *TpSil3* R | *AGCATCCTCA*TCAATGATGATGATGATGATG | 22 |
| pOESil3 pOESil1 part F | *TCATCATTGA*TGAGGATGCTCATCGTTTTG | 23 |
| pOESil3 pOESil1 part R | *AAGTCTTCAT*GATTATGATAGTTGTTGGTTCC | 24 |
| *TpSil1* F | ATGAAAGTTACCACGTCAATC | 25 |
| *TpSil1* R | TCAATTCAGAAAGAAGGAC | 26 |
| *TpSil3* F | ATGAAGACTTCTGCCATTG | 27 |
| *TpSil3* R | TCAAGCGCTCATGGAGTG | 28 |
| Blunt Simple F | CTCTGACTTGAGCGTCGA | 29 |
| pKOSil *Cas9-YFP* F | *TATCATAATC*ATGGACAAGAAGTACTCC | 30 |
| pKOSil *Cas9-YFP* R | *AGCATCCTCA*TCACTTGTACAGCTCGTC | 31 |
| pKOSil pOESil part 1 F | *ATGTGACCGA*CAGCAAAAGGCCAGGAAC | 32 |
| pKOSil pOESil part 1 R | *TCTTGTCCAT*GATTATGATAGTTGTTGGTTCCTTTG | 33 |
| pKOSil pOESil part 2 F | *GTACAAGTGA*TGAGGATGCTCATCGTTTTG | 34 |
| pKOSil pOESil part 2 R | *TTCTAGTAGC*GCCTTTTGCTCACATGTTC | 35 |
| pKOSil U6sgRNAScaffold F | *AGCAAAAGGC*GCTACTAGAATTCGAGCTCG | 36 |
| pKOSil U6sgRNAScaffold R | *CCTTTTGCTG*TCGGTCACATGTGCATCC | 37 |

Overlap regions are shown in Italic, the hexa-histidine tag is shown in Bold.

**Table S2.** Oligonucleotides for screening genetically transformed strains and qRT-PCR analysis.

| **Primer** | **Sequence** | **Primer No.** |
|---|---|---|
| OESil1 Screen F | GGAGACTACCCGCAAAAC | 1 |
| OESil1 Screen R | GAAACGATACTCGCAACG | 2 |
| OESil3 Screen F | TATGTGTATGCCCTTCGC | 3 |
| OESil3 Screen R | GTCTTGTTTCCTTCTCGC | 4 |
| KOSil1 Screen F (Band Shift) | TTTTAGCAAATCAACCATTC | 5 |
| KOSil1 Screen R (Band Shift) | GTGTTTACACCCCTCGCT | 6 |
| KOSil1 Screen F (Band Deletion) | TAATGAAAGTTACCACGTC | 7 |
| KOSil1 Screen R (Band Deletion) | CTCGTTTCTTCATCATA | 8 |
| KOSil3 Screen F (Band Shift) | CACCCTCCCTTCCTCTCCTT | 9 |
| KOSil3 Screen R (Band Shift) | GAACTCACGCTTGCTTCTCAG | 10 |
| Actin-Like/269504 F | CTCCCAATCCTGGCAATAGA | 11 |
| Actin-Like/269504 R | CGAAACCTATCCACGACGTT | 12 |
| *TpSil1/2* qRT-PCR F | TTGGTCCACCCAAAACCTCC | 13 |
| *TpSil1/2* qRT-PCR F | GAGTGTGGGAGATTCCGTCG | 14 |
| *TpSil3* qRT-PCR F | GTGCAAAGAGTGCCAAGATGAG | 15 |
| *TpSil3* qRT-PCR R | CTTGTGTGACTTGGCCATGC | 16 |

**Table S3.** Oligonucleotides for encoding sgRNAs.

| Name | Sequence |
|---|---|
| *TpSil1*_sgRNA1 | CGGCGCTGCAGATGTGCAGA **GGG** |
| *TpSil1*_sgRNA2 | TCTACTATGATGAAGAAACG **AGG** |
| *TpSil1*_sgRNA3 | TGCAGTAATGAGGCCACCGG **TGG** |
| *TpSil3*_sgRNA1 | TCAAGAGTTCTCAATCGGCG **GGG** |
| *TpSil3*_sgRNA2 | TTTCGAGCTGTGCATGGACA **TGG** |
| *TpSil3*_sgRNA3 | GAGAACTCTTGAAGGACATG **GGG** |

Protospacer adjacent motif (PAM) is shown in Bold.

**Table S4.** Incubation system for Cas9 and sgRNA-mediated *in vitro* DNA cleavage.

| Component | Volume (μL) |
| --- | --- |
| sgRNA (6 pM) | 0.6 |
| Cas9 Nuclease (5 pM, Vazyme, EN301) | 5 |
| Cas9 Nuclease Buffer (10×) | 2 |
| RNase-Free $H_2O$ | 8.4 |
| Total Volume | 16 |

**Table S5.** Off-target prediction for up to five mismatches.

| Mismatches | Number of targets | | | |
|---|---|---|---|---|
| | *TpSil1*_sgRNA1 | *TpSil1*_sgRNA2 | *TpSil3*_sgRNA1 | *TpSil3*_sgRNA2 |
| 0 | 1* | 1 | 1 | 1 |
| 1 | 0 | 0 | 0 | 0 |
| 2 | 0 | 0 | 0 | 0 |
| 3 | 0 | 0 | 0 | 0 |
| 4 | 1 | 1 | 0 | 0 |
| 5 | 15 | 32 | 10 | 12 |

*In fact, *TpSil1*_sgRNA1 also targets *TpSil2* due to their high sequence homology. However, off-target analysis did not identify distinct genes for *TpSil2*, as only one corresponding gene, THAPSDRAFT_11360, is listed in the *Thalassiosira pseudonana* genome database.
